# Computationally designed mRNA-launched protein nanoparticle vaccines

**DOI:** 10.1101/2024.07.22.604655

**Authors:** Grace G. Hendricks, Lilit Grigoryan, Mary Jane Navarro, Nicholas J. Catanzaro, Miranda L. Hubbard, John M. Powers, Melissa Mattocks, Catherine Treichel, Alexandra C. Walls, Jimin Lee, Daniel Ellis, Jing Yang (John) Wang, Suna Cheng, Marcos C. Miranda, Adian Valdez, Cara W. Chao, Sidney Chan, Christine Men, Max R. Johnson, Harold Hui, Sheng-Yang Wu, Victor Lujan, Hiromi Muramatsu, Paulo J.C. Lin, Molly M.H. Sung, Ying K. Tam, Elizabeth M. Leaf, Norbert Pardi, Ralph S. Baric, Bali Pulendran, David Veesler, Alexandra Schäfer, Neil P. King

## Abstract

Both protein nanoparticle and mRNA vaccines were clinically de-risked during the COVID-19 pandemic^1–6^. These vaccine modalities have complementary strengths: antigen display on protein nanoparticles can enhance the magnitude, quality, and durability of antibody responses^7–10^, while mRNA vaccines can be rapidly manufactured^11^ and elicit antigen-specific CD4 and CD8 T cells^12,13^. Here we leverage a computationally designed icosahedral protein nanoparticle that was redesigned for optimal secretion from eukaryotic cells^14^ to develop an mRNA-launched nanoparticle vaccine for SARS-CoV-2. The nanoparticle, which displays 60 copies of a stabilized variant of the Wuhan-Hu-1 Spike receptor binding domain (RBD)^15^, formed monodisperse, antigenically intact assemblies upon secretion from transfected cells. An mRNA vaccine encoding the secreted RBD nanoparticle elicited 5- to 28-fold higher levels of neutralizing antibodies than an mRNA vaccine encoding membrane-anchored Spike, induced higher levels of CD8 T cells than the same immunogen when delivered as an adjuvanted protein nanoparticle, and protected mice from vaccine-matched and -mismatched SARS-CoV-2 challenge. Our data establish that delivering protein nanoparticle immunogens via mRNA vaccines can combine the benefits of each modality and, more broadly, highlight the utility of computational protein design in genetic immunization strategies.

## Introduction

The emergence of SARS-CoV-2 in late 2019^16^ and the subsequent COVID-19 pandemic highlighted the need for ultrapotent and rapidly scalable vaccine platforms^11,17,18^. Lipid nanoparticle (LNP)-encapsulated, nucleoside-modified mRNA vaccines encoding prefusion-stabilized, membrane-anchored Spike (S-2P) were found to be safe, effective, and manufacturable at scale, leading to emergency use authorization less than a year after the sequence of the viral genome was available^3,5,19–22^. These first-generation mRNA vaccines saved many lives and lessened the global health and economic burden of SARS-CoV-2^23^. Although mRNA-LNPs were not the only vaccine modality utilized in response to the COVID-19 pandemic^24^, the subsequent introduction of several booster vaccines to keep up with emerging immune-evasive viral variants^25^ has further emphasized the sequence-invariant manufacturing advantages of mRNA vaccines^26,27^.

We previously described an *in vitro*-assembled protein nanoparticle vaccine displaying 60 copies of the Wuhan-Hu-1 SARS-CoV-2 Spike receptor binding domain (RBD) on the computationally designed two-component icosahedral nanoparticle I53-50^28^. The resultant nanoparticle immunogen, RBD-I53-50, elicited robust neutralizing antibody responses in mice and non-human primates^29,30^. Furthermore, RBD-I53-50 was found to be safe and immunogenic in clinical trials^2,31^, prompting its licensure in multiple jurisdictions under the name SKYCovione™. Consistent with previous studies indicating that multivalent antigen display improves the magnitude, breadth, and durability of vaccine-elicited immune responses^7–10^, three doses of RBD-I53-50 not only protected against heterologous Omicron BA.1 challenge in non-human primates, but also elicited broadly neutralizing antibodies against other sarbecoviruses^32,33^.

Clinical de-risking of these technologies during the COVID-19 pandemic has motivated further technological development of both the mRNA-LNP and protein nanoparticle vaccine modalities. In particular, there has been a push to combine the potency of protein nanoparticle immunogens with the speed of mRNA vaccine manufacture^34–37^. To successfully develop this platform, protein nanoparticle immunogens must be designed such that they are not only produced and assembled within eukaryotic host cells, but also efficiently secreted. To this end, we recently developed a general computational method that improves the secretion of designed protein nanoparticles without perturbing self-assembly^14^. However, the performance of these computationally designed, secretion-optimized protein nanoparticles as genetically encoded vaccines is only beginning to be characterized^38^.

Here, we develop and demonstrate proof-of-concept for computationally designed mRNA-launched protein nanoparticle vaccines. We found that an mRNA vaccine encoding a secreted RBD nanoparticle elicited more potent, broad, and protective antibody responses than mRNA vaccines encoding prefusion-stabilized, membrane-anchored Spike or secreted trimeric RBD, demonstrating the superiority of particulate immunogens even in the context of genetic immunization.

## Results

### Immunogen design and characterization

To generate a secreted RBD nanoparticle vaccine candidate, we multivalently displayed the Wuhan-Hu-1 SARS-CoV-2 Spike RBD on the exterior surface of the self-assembling protein nanoparticle I3-01NS^14,39^. I3-01NS is a one-component, 60-subunit complex with icosahedral symmetry derived from a naturally occurring bacterial aldolase^40^ that was computationally redesigned for optimal secretion from mammalian cells. We genetically fused the RBD (residues 328–531) to the N terminus of I3-01NS using a 16-residue glycine/serine linker to enable flexible presentation of the antigen extending from the nanoparticle surface (**Fig. 1a**). The resultant fusion construct, RBD-I3-01NS, was recombinantly expressed in human (Expi293F) cells to mimic the process of expression and secretion during genetic immunization. SDS-PAGE of the cell culture supernatant revealed that RBD-I3-01NS did not secrete (**Extended Data Fig. 1a**). To recover secretion, we swapped out the wild-type Wuhan-Hu-1 SARS-CoV-2 RBD for a stabilized and higher-yielding version, Rpk9, which contains three mutations (Y365F, F392W, V395I) that repack the linoleic acid binding pocket^15,41,42^. The new fusion construct, Rpk9-I3-01NS, did secrete from cells and was carried forward for purification (**Extended Data Fig. 1a**). Size exclusion chromatography (SEC) of Rpk9-I3-01NS revealed a predominant peak corresponding to the target icosahedral assembly (**Fig. 1b**), and dynamic light scattering (DLS) and negative stain electron microscopy (nsEM) confirmed a homogenous and monodisperse population of nanoparticles **Fig. 1c, d**). Biolayer interferometry (BLI) with Fc-tagged ACE2 (hACE2-Fc), the class 4 RBD-directed monoclonal antibody (mAb) CR3022^43,44^, and the class 3 RBD-directed mAb S309^44,45^ confirmed that multiple epitopes of Rpk9 were intact and accessible in the context of the I3-01NS nanoparticle (**Fig. 1e**). To rigorously assess the yield of secreted Rpk9-I3-01NS, we used the purified protein to create a standard curve for supernatant enzyme-linked immunosorbent assays (ELISAs). Based on three independent transfections, Rpk9-I3-01NS was shown to secrete at ∼30 mg/L whereas the original RBD-I3-01NS construct secreted at levels below the lower limit of detection of the assay (**Extended Data Fig. 1b-d**). These data indicate that Rpk9-I3-01NS nanoparticles secrete efficiently and are biochemically, antigenically, and structurally intact.

**Fig. 1.**
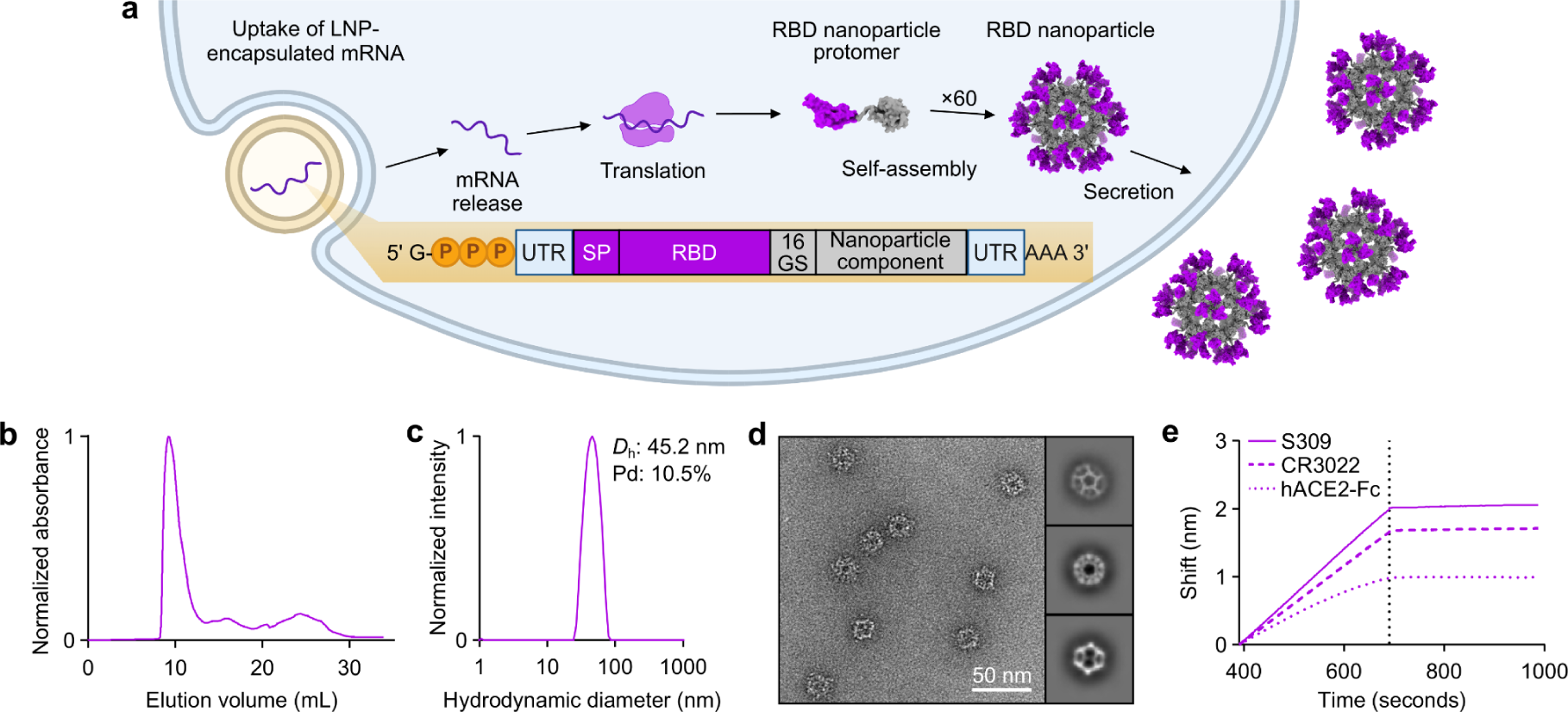
Design and characterization of a secretable SARS-CoV-2 RBD nanoparticle. **a,** Schematic of the biogenesis of secreted RBD nanoparticles using LNP-encapsulated mRNA as an example for method of delivery. The secretory pathway has been omitted for simplicity. UTR, untranslated region; SP, signal peptide; 16 GS, 16-residue glycine/serine linker. The protein models and schematic were rendered using ChimeraX^46^ and BioRender.com, respectively. **b,** Size exclusion chromatogram of Rpk9-I3-01NS purification. **c,** Dynamic light scattering of SEC-purified Rpk9-I3-01NS. *D*_h_, hydrodynamic diameter; Pd, polydispersity. **d,** Representative electron micrograph of negatively stained SEC-purified Rpk9-I3-01NS and 2D class averages. **e,** Binding of immobilized hACE2-Fc, CR3022, and S309 to SEC-purified Rpk9-I3-01NS as assessed by biolayer interferometry. The dotted vertical line separates the association and dissociation steps.

### Elicitation of neutralizing antibody responses

We assessed the immunogenicity of Rpk9-I3-01NS when delivered as an adjuvanted protein vaccine (“protein-delivered”) and an mRNA vaccine (“mRNA-launched”). To directly evaluate the impact of nanoparticle formation, we also assessed the immunogenicity of protein-delivered and mRNA-delivered “non-assembling” Rpk9-I53-50A trimers. I53-50A is a trimeric scaffold that shares 91% amino acid sequence identity with I3-01NS but lacks the hydrophobic interface that drives nanoparticle assembly. We also included the protein-delivered S-2P-foldon trimer as a benchmark immunogen. Finally, to assess Rpk9-I3-01NS within the current COVID-19 vaccine landscape, we also evaluated the immunogenicity of mRNA-delivered membrane-anchored S-2P trimers (using the exact mRNA sequence of Comirnaty^®^) as well as protein-delivered Rpk9-I53-50 two-component nanoparticles, similar to RBD-I53-50 (licensed as SKYCovione™). Groups of ten BALB/c mice were immunized intramuscularly on weeks 0 and 3 with either AddaVax-adjuvanted protein (equimolar amounts of RBD: 0.9 μg of RBD per dose for Rpk9-based constructs, 5 μg of Spike per dose for S-2P-foldon) or nucleoside-modified, LNP-encapsulated mRNA (0.2, 1, or 5 μg dose) (**Fig. 2a and Supplementary Table 1, 2**). Additionally, we immunized five mice with lipid nanoparticles not encapsulating any mRNA (“empty” LNPs) as a negative control group. Vaccine-elicited antibody responses were then assessed two weeks post-prime and -boost via serum ELISAs and pseudovirus neutralization assays.

**Fig. 2.**
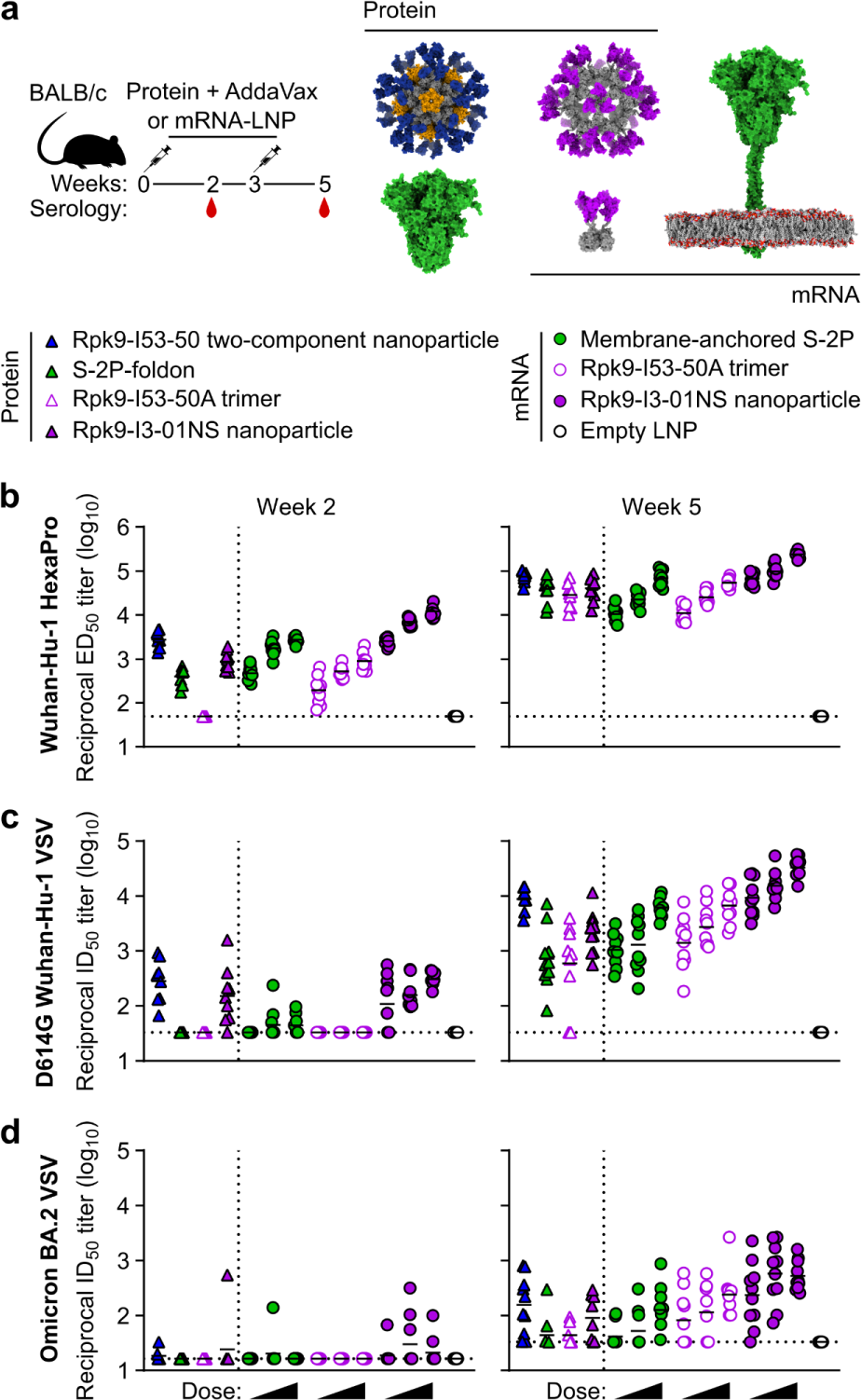
mRNA-launched RBD nanoparticles elicit potent neutralizing antibody titers in BALB/c mice. **a,** Study design and groups; n=10 mice/group received either AddaVax-adjuvanted protein (equimolar amounts of RBD: 0.9 μg of RBD per dose for Rpk9-based constructs, 5 μg of Spike per dose for S-2P-foldon) or nucleoside-modified, LNP-encapsulated mRNA (0.2, 1, or 5 µg per dose). Molecular models of immunogens are not to scale. Rpk9 shown in blue (Rpk9-I53-50) or purple (Rpk9-I53-50A and Rpk9-I3-01NS). Trimer scaffold shown in gray (Rpk9-I53-50, Rpk9-I53-50A, and Rpk9-I3-01NS). Pentamer component shown in orange (Rpk9-I53-50). S-2P-foldon and membrane-anchored S-2P, shown in green, are adapted from ref. 51. Glycans have been omitted from all models for simplicity. All models were rendered using ChimeraX^46^. **b,** Serum antibody binding titers against Wuhan-Hu-1 SARS-CoV-2 S HexaPro^52^, measured by ELISA. **c,** Serum neutralizing antibody titers against VSV pseudotyped with D614G Wuhan-Hu-1 SARS-CoV-2 S. **d,** Serum neutralizing antibody titers against VSV pseudotyped with Omicron BA.2 SARS-CoV-2 S. **b-d,** Each symbol represents an individual animal and the geometric mean titer (GMT) from each group is indicated by a horizontal line. The dotted horizontal line represents the lowest limit of detection for the assay; limits of detection vary between groups. The dotted vertical line separates the protein and mRNA immunized groups. Statistical analyses have been omitted for clarity but can be found in **Supplementary Information.**

Two weeks post-prime, all RBD nanoparticle vaccines consistently elicited antigen-specific binding and vaccine-matched (D614G Wuhan-Hu-1 VSV) neutralizing antibody titers. By contrast, the non-particulate vaccines consistently elicited antigen-specific binding antibody titers but minimal to no vaccine-matched neutralizing antibody titers (**Fig. 2b, c and Extended Data Fig. 2, 3**). Unsurprisingly, minimal to no vaccine-mismatched (Omicron BA.2 VSV) neutralizing antibody titers were elicited by any of the vaccines (**Fig. 2d and Extended Data Fig. 4**). Notably, at every dose of mRNA, mRNA-launched Rpk9-I3-01NS elicited ∼4-fold higher antigen-specific binding antibody titers than mRNA-delivered membrane-anchored S-2P. Furthermore, at every dose of mRNA, mRNA-launched Rpk9-I3-01NS elicited >10-fold higher antigen-specific binding antibody titers than secreted Rpk9-I53-50A, despite secreting at ∼4-fold lower levels in tissue culture (**Extended Data Fig. 1**). These observations were consistent with the intrinsically higher immunogenicity of the particulate immunogens when delivered as antigen dose-matched adjuvanted proteins: Rpk9-I3-01NS and Rpk9-I53-50 both elicited >15-fold higher antigen-specific binding antibody titers than Rpk9-I53-50A and ∼2-fold and ∼7-fold higher than S-2P-foldon, respectively.

Following a second immunization, we observed enhanced antigen-specific binding, vaccine-matched neutralizing, and vaccine-mismatched neutralizing antibody titers for all vaccines (**Fig. 2b-d and Extended Data Fig. 2**-4). Consistent with post-prime data, mRNA-launched Rpk9-I3-01NS elicited ≥5-fold higher vaccine-matched and ≥4-fold higher vaccine-mismatched neutralizing antibody titers than mRNA-delivered membrane-anchored S-2P at every dose of mRNA. Additionally, mRNA-launched Rpk9-I3-01NS elicited ≥5-fold higher vaccine-matched and ≥2-fold higher vaccine-mismatched neutralizing antibody titers than secreted Rpk9-I53-50A. Notably, the level of vaccine-matched and -mismatched neutralizing antibody titers elicited by the 0.2 μg dose of mRNA-launched Rpk9-I3-01NS were comparable to those elicited by the 5 μg dose of mRNA-delivered membrane-anchored S-2P, despite a 25-fold lower dose of mRNA. These observations were consistent with post-prime data and were further corroborated when the immunogens were delivered as antigen dose-matched adjuvanted proteins: Rpk9-I3-01NS elicited ∼3-fold higher vaccine-matched and ∼2-fold higher vaccine-mismatched neutralizing antibody titers than both Rpk9-I53-50A and S-2P-foldon, while Rpk9-I53-50 elicited >10-fold higher vaccine-matched and ∼3-fold higher vaccine-mismatched neutralizing antibody titers. Interestingly, when comparing vaccine modalities, the 0.2 μg dose of mRNA-launched Rpk9-I3-01NS elicited comparable vaccine-matched and -mismatched neutralizing antibody titers to the 0.9 μg RBD dose of protein-delivered two-component Rpk9-I53-50. Finally, as expected, we also detected antibody responses to the I3-01NS, I53-50A, and I53-50 scaffolds (**Extended Data Fig. 5**).

Several conclusions can be drawn from these data. First, secreted RBD nanoparticles can be effectively delivered as mRNA vaccines and elicit potent humoral responses. Second, as shown previously^1,15,28,47–50^, multivalent display of RBDs on self-assembling protein nanoparticles improves immunogenicity, as indicated by the enhanced immunogenicity of the adjuvanted protein RBD nanoparticles compared to non-assembling RBD trimers at equivalent doses of RBD. Third, mRNA-launched RBD nanoparticles likely assemble properly *in vivo*, as indicated by the enhanced immunogenicity of the mRNA-launched RBD nanoparticles compared to non-assembling RBD trimers at equivalent doses of mRNA. Fourth, mRNA-launched RBD nanoparticles are as, if not more, immunogenic than adjuvanted two-component RBD nanoparticle protein vaccines, depending on the dose of mRNA. Finally, mRNA-launched RBD nanoparticles are several-fold more immunogenic than COVID-19 mRNA vaccines encoding membrane-anchored S-2P or a secreted RBD trimer. In summary, multivalent antigen display on computationally designed protein nanoparticles enhances immunogenicity, whether delivered as adjuvanted protein or mRNA vaccines.

### Induction of antigen-specific T cell responses

We next evaluated the antigen-specific T cell response induced by Rpk9-I3-01NS when delivered as an adjuvanted protein vaccine and an mRNA vaccine. As before, we also evaluated protein-delivered S-2P-foldon trimers and Rpk9-I53-50A trimers, as well as mRNA-delivered membrane-anchored S-2P trimers and secreted Rpk9-I53-50A trimers. Groups of five C57BL/6 mice were immunized intramuscularly at weeks 0 and 3 with either AddaVax-adjuvanted protein (equimolar amounts of RBD: 0.9 μg of RBD per dose for Rpk9-based constructs, 5 μg of Spike per dose for S-2P-foldon) or nucleoside-modified, LNP-encapsulated mRNA (1 µg dose) (**Fig. 3a**). Three weeks post-boost, lung and spleen lymphocytes were isolated and stimulated *in vitro* with a peptide pool containing overlapping peptides from the Wuhan-Hu-1 SARS-CoV-2 Spike protein. Following stimulation, intracellular staining was performed to detect the production of individual cytokines (**Extended Data Fig. 6**). For a baseline, we also isolated and stimulated lung and spleen lymphocytes from naive (i.e., unimmunized) mice. As expected, we observed minimal to undetectable antigen-specific CD4 or CD8 T cells in either the lungs or spleens of these mice (**Fig. 3b, c**).

**Fig. 3.**
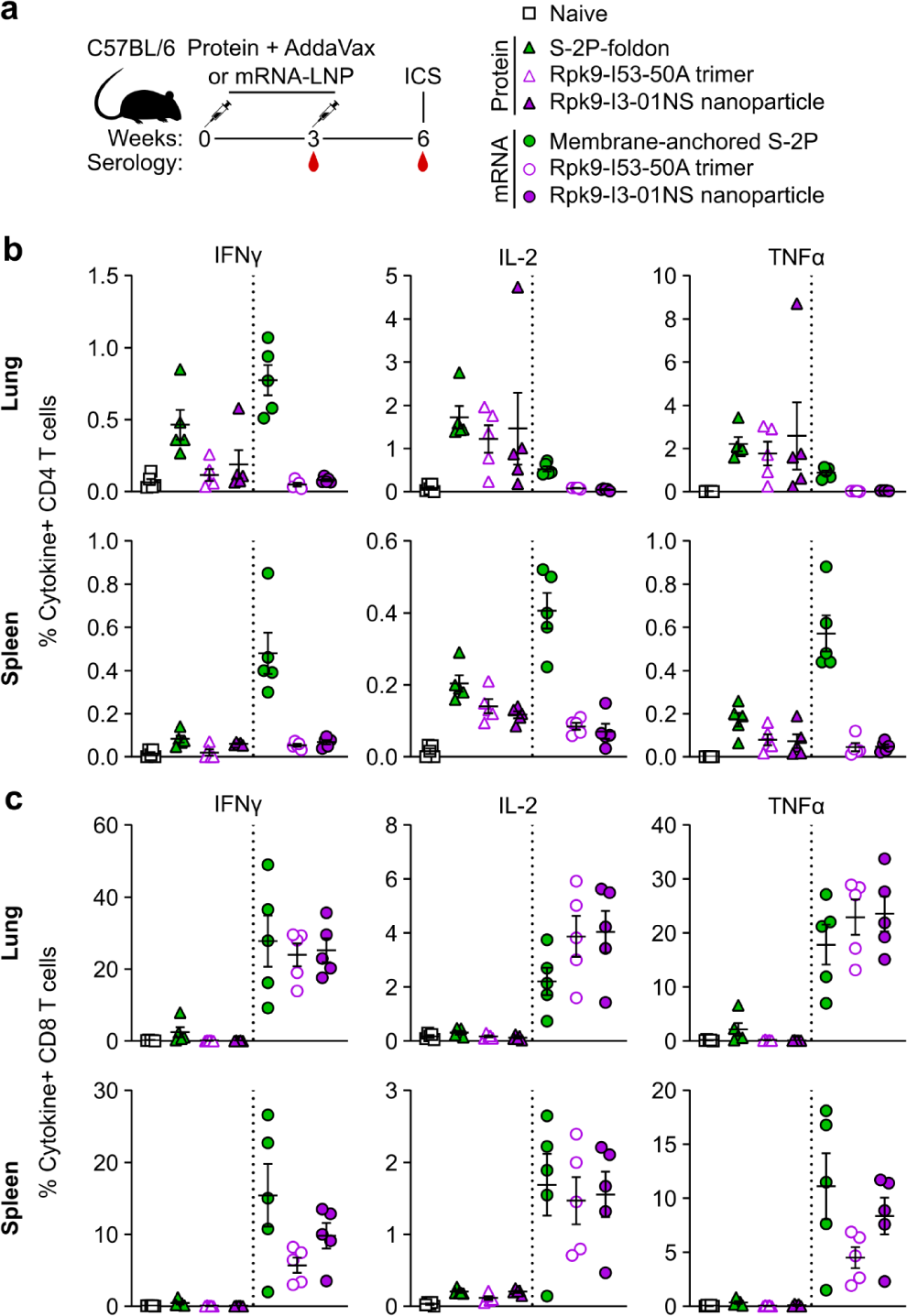
mRNA vaccines induce robust antigen-specific CD8 T cell responses in C57BL/6 mice. **a,** Study design and groups (ICS, intracellular cytokine staining); n=10 mice/group received either AddaVax-adjuvanted protein (equimolar amounts of RBD: 0.9 μg of RBD per dose for Rpk9-based constructs, 5 μg of Spike per dose for S-2P-foldon) or nucleoside-modified, LNP-encapsulated mRNA (1 µg per dose). **b,** Percentage of CD4 T cells producing IFNγ, IL-2, and TNFα in response to a Wuhan-Hu-1 SARS-CoV-2 S peptide pool of overlapping 15-mers. **c,** Percentage of CD8 T cells producing IFNγ, IL-2, and TNFα in response to a Wuhan-Hu-1 SARS-CoV-2 S peptide pool of overlapping 15-mers. **b-c,** Each symbol represents an individual animal. Error bars represent mean ± SEM. The dotted vertical line separates the protein and mRNA immunized groups. Statistical analyses have been omitted for clarity but can be found in **Supplementary Information.**

With regard to antigen-specific CD4 T cell responses, mRNA-delivered membrane-anchored S-2P induced the highest levels of IFNγ production in the lungs (∼0.8% IFNγ+ CD4 T cells), although higher levels of IL-2 and TNFα production were observed with the protein-delivered vaccines (∼1.2–1.7% IL-2+ CD4 T cells and ∼1.8–2.6% TNFα+ CD4 T cells) (**Fig. 3b**). In the spleen, mRNA-delivered membrane-anchored S-2P induced the highest levels of IFNγ, IL-2, and TNFα production among all groups (∼0.5% IFNγ+, ∼0.4% IL-2+, and ∼0.6% TNFα+ CD4 T cells). In both the lungs and spleen, we also observed that protein-delivered S-2P-foldon induced slightly higher levels of IFNγ, IL-2, and TNFα production compared to the other adjuvanted protein vaccines, although the differences were not statistically significant. We attribute these observations in part to the fact that the Spike-based immunogens comprise all of the possible T cell epitopes in the overlapping peptide library used for stimulation, whereas the RBD-based immunogens comprise only a fraction of the epitopes^53,54^. With regard to antigen-specific CD8 T cell responses, all three mRNA vaccines induced comparable levels of IFNγ, IL-2, and TNFα production in the lungs (∼24–28% IFNγ+, ∼2–4% IL-2+, and ∼18–24% TNFα+ CD8 T cells), as well as comparable IL-2 production in the spleen (∼1.5–1.6% IL-2+ CD8 T cells) (**Fig. 3c**). However, mRNA-delivered membrane-anchored S-2P induced higher levels of IFNγ and TNFα production in the spleen (∼16% IFNγ+ and ∼11% TNFα+ CD8 T cells) compared to all other groups, although the differences were not always statistically significant. As expected, all three adjuvanted protein vaccines induced minimal or undetectable antigen-specific CD8 T cell responses in either the lungs or spleen.

We note that our use of a peptide pool spanning the Spike, rather than a pool matched to each full-length immunogen, means we have likely not captured the whole T cell response induced by the RBD nanoparticles and RBD trimers, as we have previously shown that computationally designed nanoparticle scaffolds themselves contain T cell epitopes^30^. Nonetheless, these results still corroborate previous studies in showing that mRNA vaccines induce significantly more robust CD8 T cell responses when compared to protein-delivered vaccines, even in the presence of adjuvants^55^. We also note that the high frequencies of CD8 T cells elicited by the mRNA vaccines studied here are similar to those elicited by licensed COVID-19 vaccines in previous studies in mice^13^ but that the frequencies observed in humans from the same vaccines were considerably lower^12^.

In addition to evaluating the T cell response, we also performed serological analyses (immunized groups only) three weeks post-prime and -boost to evaluate immunogenicity in a second mouse model (**Extended Data Fig. 7, 8**). The results were generally consistent with those observed in BALB/c mice (**Fig. 2b, c and Extended Data Fig. 2, 3**). However, in C57BL/6 mice, there appeared to be a smaller enhancement of immunogenicity provided by the mRNA-launched RBD nanoparticles compared to mRNA-delivered membrane-anchored S-2P.

### Protection against challenge with mouse-adapted SARS-CoV-2

We next evaluated the protective efficacy of mRNA-launched Rpk9-I3-01NS nanoparticles compared to mRNA-delivered membrane-anchored S-2P trimers and secreted Rpk9-I53-50A trimers. We also evaluated an mRNA vaccine encoding luciferase as a control. BALB/c mice were immunized intramuscularly with either nucleoside-modified, LNP-encapsulated mRNA (1 µg dose) or an equivalent volume of phosphate-buffered saline (PBS) (**Fig. 4a**). Five weeks after the single immunization, vaccine-elicited antibody responses were then assessed via serum neutralization assays with vaccine-matched (D614G Wuhan-Hu-1 SARS-CoV-2) authentic virus. Notably, mRNA-launched Rpk9-I3-01NS elicited ∼28-fold and ∼11-fold higher vaccine-matched neutralizing antibody titers than mRNA-delivered membrane-anchored S-2P and secreted Rpk9-I53-50A, respectively (**Fig. 4b and Extended Data Fig. 9**). As expected, immunization with PBS and luciferase mRNA elicited little to no vaccine-matched neutralization. One week later (i.e., six weeks after the single immunization), the mice were challenged intranasally with 1×10^5^ plaque-forming units (PFUs) of mouse-adapted Wuhan-Hu-1 SARS-CoV-2 MA10^56^ and followed for four days to assess protection from disease. All mice immunized with mRNA vaccines encoding SARS-CoV-2 immunogens were protected against weight loss and death for four days post infection (**Fig. 4c**). In contrast, mice immunized with PBS or mRNA encoding luciferase experienced weight loss up to 20% of their starting body weight. Additionally, one PBS-immunized mouse and one luciferase mRNA-immunized mouse had succumbed to infection by the fourth day. Consistent with these data, we observed severe lung discoloration in PBS and luciferase mRNA-immunized mice four days post infection, an indication of severe disease (**Fig. 4d**). By contrast, mice immunized with mRNA encoding SARS-CoV-2 immunogens showed minimal to no lung discoloration four days post infection.

**Fig. 4.**
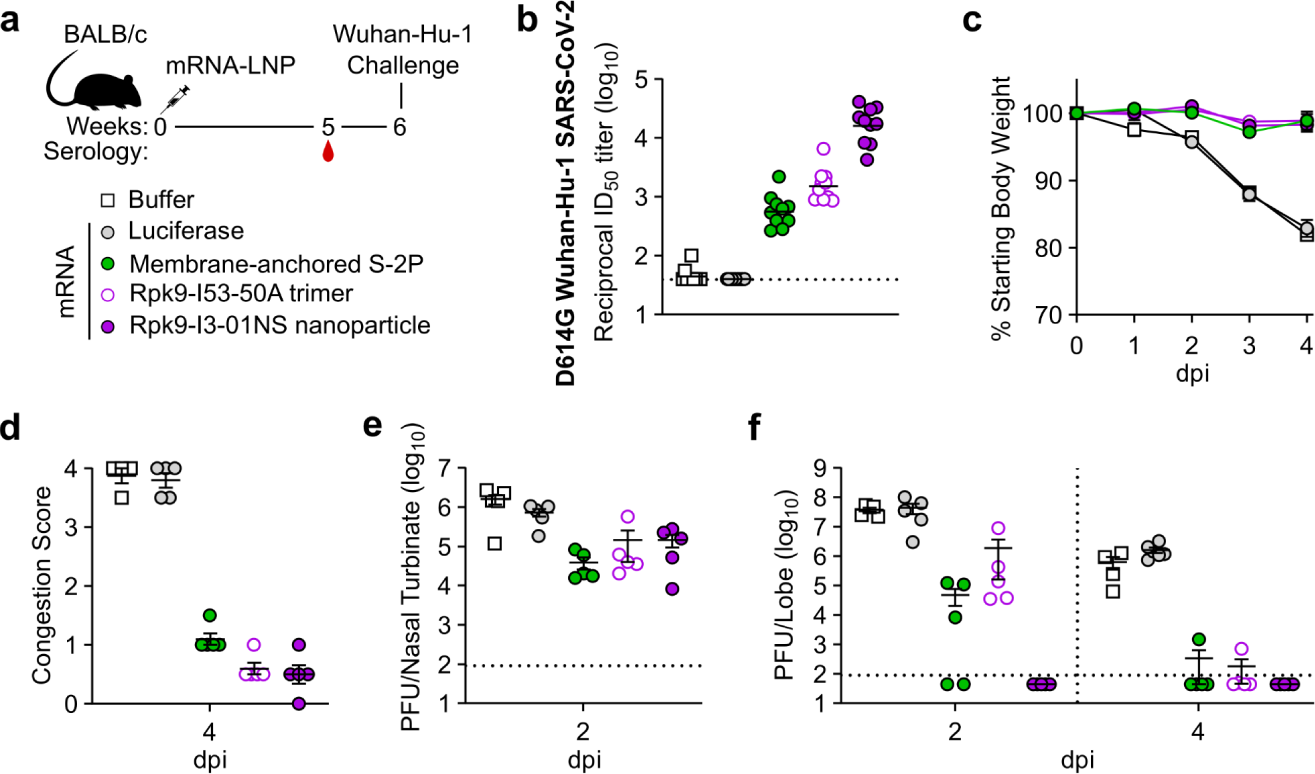
mRNA vaccines confer protective immunity against mouse-adapted SARS-CoV-2. **a,** Study design and groups; n=4-6 mice/group/time point (2 and 4 days post infection (dpi)) received either nucleoside-modified, LNP-encapsulated mRNA (1 µg dose) or an equivalent volume of phosphate-buffered saline (PBS). **b,** Serum neutralizing antibody titers against D614G Wuhan-Hu-1 SARS-CoV-2 authentic virus. Each symbol represents an individual animal and the GMT from each group is indicated by a horizontal line. The dotted horizontal line represents the lowest limit of detection for the assay. **c,** Weight loss up to 4 days dpi. Each symbol is the mean of the group for the time point ± SEM. **d,** Congestion score at 4 dpi (scored as: 0 = no discoloration, 4 = severe discoloration). **e,** Infectious viral load at 2 dpi in the nasal cavity after challenge of vaccinated mice as determined by plaque assay. **f,** Infectious viral load at 2 and 4 dpi in the lung after challenge of vaccinated mice as determined by plaque assay. The dotted vertical line separates time points. The dotted horizontal line indicates the limit of detection; for samples with values below this, data are plotted at half the limit of detection. **b-f,** Statistical analyses have been omitted for clarity but can be found in **Supplementary Information. d-f,** Each symbol represents an individual animal. Error bars represent mean ± SEM.

Analysis of viral titers in nasal turbinates and lung tissue further elucidated the performance of the mRNA vaccines in the upper and lower respiratory tract, respectively. Two days post infection, we observed moderate viral loads in nasal turbinates across all groups (3.9×10^4^–1.6×10^6^ PFU/tissue), albeit at slightly lower levels in mice immunized with mRNA encoding SARS-CoV-2 immunogens (**Fig. 4e**). These results are consistent with previous reports suggesting that prevention of viral replication in the upper respiratory tract can be difficult to achieve by intramuscular vaccination^57,58^. Nonetheless, two days post infection, when viral replication of mouse-adapted Wuhan-Hu-1 SARS-CoV-2 MA10 has been shown to reach its peak in the lung^56^, mRNA-launched Rpk9-I3-01NS provided complete protection from detectable virus in the lung (**Fig. 4f**). By contrast, mice immunized with mRNA-delivered membrane-anchored S-2P and secreted Rpk9-I53-50A had moderate viral loads in the lung (4.9×10^4^ and 1.9×10^6^ PFU/tissue, respectively), and mice immunized with PBS or luciferase mRNA had high viral loads (3.9×10^7^ and 4.4×10^7^ PFU/tissue, respectively). These differences in lung viral titers at two days post infection are consistent with our authentic virus neutralization data, and align with previous reports showing that antigen-specific binding and vaccine-matched neutralizing antibody titers are correlates of protection against SARS-CoV-2^59–62^. Four days post infection, we observed a decrease in lung viral titers across all groups. As expected, all five mice immunized with mRNA-launched Rpk9-I3-01NS maintained undetectable viral loads in the lung. In addition, most mice immunized with mRNA-delivered membrane-anchored S-2P and secreted Rpk9-I53-50A had undetectable viral loads at this time point, and the mice immunized with PBS or luciferase mRNA had moderate viral loads (5.4×10^5^ and 1.6×10^6^ PFU/tissue, respectively). Altogether, these results indicate that mRNA-launched RBD nanoparticles elicit more protective antibodies and provide better protection against viral replication in the lung during vaccine-matched infection than COVID-19 mRNA vaccines encoding membrane-anchored S-2P or a secreted RBD trimer.

### Protection against challenge with mouse-adapted Omicron BA.5 SARS-CoV-2

Finally, we evaluated the protective efficacy of mRNA-launched Rpk9-I3-01NS nanoparticles against vaccine-mismatched challenge compared to mRNA-delivered membrane-anchored S-2P trimers and secreted Rpk9-I53-50A trimers. We again included an mRNA vaccine encoding luciferase as a control. BALB/c mice were immunized intramuscularly at weeks 0 and 4 with either nucleoside-modified, LNP-encapsulated mRNA (1 or 5 µg dose) or an equivalent volume of phosphate-buffered saline (PBS) (**Fig. 5a**). Vaccine-elicited antibody responses were assessed four weeks post-boost immunization via serum neutralization assays with vaccine-mismatched (Omicron BA.5 SARS-CoV-2) authentic virus. Notably, for both doses of mRNA, mRNA-launched Rpk9-I3-01NS elicited ≥5-fold higher vaccine-mismatched neutralizing antibody titers than both mRNA-delivered membrane-anchored S-2P and secreted Rpk9-I53-50A (**Fig. 5b and Extended Data Fig. 10**). As expected, immunization with PBS and luciferase mRNA elicited little to no vaccine-matched neutralization. One week later (i.e., five weeks post-boost), the mice were challenged intranasally with 1×10^5^ plaque-forming units (PFUs) of mouse-adapted Omicron BA.5 SARS-CoV-2^63^ and followed for four days to assess protection from disease. All mice immunized with mRNA vaccines encoding SARS-CoV-2 immunogens were protected against weight loss and death for four days post infection, irrespective of dose of mRNA (**Fig. 5c**). In contrast, mice immunized with PBS or mRNA encoding luciferase experienced weight loss up to 20% of their starting body weight. Additionally, one PBS-immunized mouse and one luciferase mRNA-immunized mouse (from the 1 µg dose group) had succumbed to infection by the fourth day. Consistent with these data, we observed severe lung discoloration in PBS and luciferase mRNA-immunized mice four days post infection, indicative of inflammation, edema, and diffuse alveolar damage (**Fig. 5d**). By contrast, mice immunized with mRNA encoding SARS-CoV-2 immunogens showed minimal lung discoloration four days post infection.

**Fig. 5.**
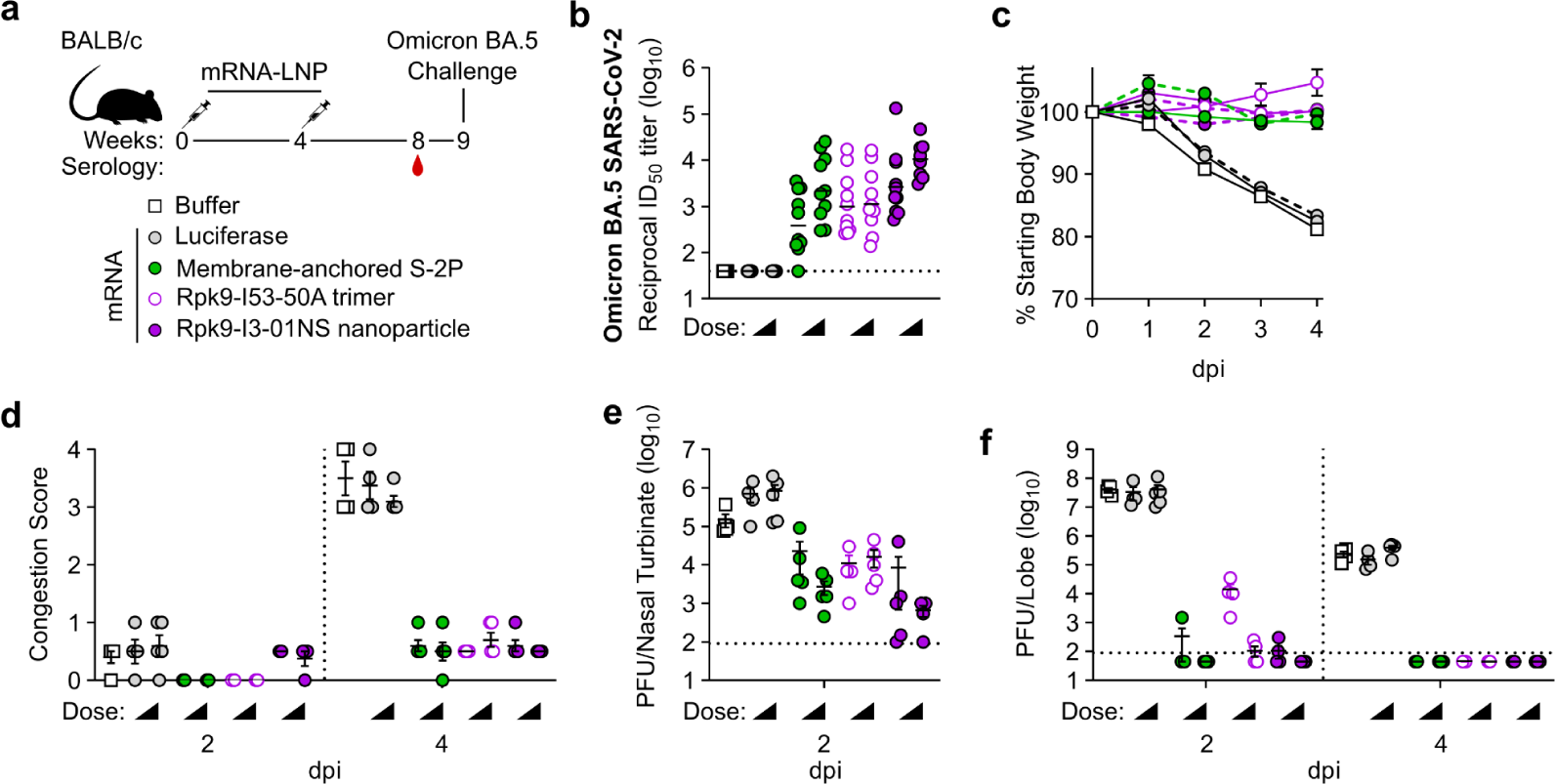
mRNA vaccines confer protective immunity against mouse-adapted Omicron BA.5 SARS-CoV-2. **a,** Study design and groups; n=4-5 mice/group/time point (2 and 4 days post infection (dpi)) received either nucleoside-modified, LNP-encapsulated mRNA (1 or 5 µg dose) or an equivalent volume of phosphate-buffered saline (PBS). **b,** Serum neutralizing antibody titers against Omicron BA.5 SARS-CoV-2 authentic virus. Each symbol represents an individual animal and the GMT from each group is indicated by a horizontal line. The dotted horizontal line represents the lowest limit of detection for the assay. **c,** Weight loss up to 4 dpi. Each symbol is the mean of the group for the time point ± SEM. The solid lines correspond to groups immunized with 1 µg of mRNA; the dashed lines correspond to groups immunized with 5 µg of mRNA. **d,** Congestion score at 2 and 4 dpi (scored as: 0 = no discoloration, 4 = severe discoloration). The dotted vertical line separates time points. **e,** Infectious viral load at 2 dpi in the nasal cavity after challenge of vaccinated mice as determined by plaque assay. The dotted horizontal line indicates the limit of detection. **f,** Infectious viral load at 2 and 4 dpi in the lung after challenge of vaccinated mice as determined by plaque assay. The dotted vertical line separates time points. The dotted horizontal line indicates the limit of detection; for samples with values below this, data are plotted at half the limit of detection. **b-f,** Statistical analyses have been omitted for clarity but can be found in **Supplementary Information. d-f,** Each symbol represents an individual animal. Error bars represent mean ± SEM.

Analysis of viral titers in nasal turbinates and lung tissue further elucidated the performance of the mRNA vaccines in the upper and lower respiratory tract, respectively. Two days post infection, we observed low viral loads in nasal turbinates across all groups immunized with mRNA encoding SARS-CoV-2 immunogens (6.6×10^2^–2.3×10^4^ PFU/tissue), irrespective of mRNA dose (**Fig. 5e**). Notably, two mice immunized with mRNA-launched Rpk9-I3-01NS (one per dose of mRNA) were completely protected from detectable virus in the nasal turbinates. In contrast, mice immunized with PBS or luciferase mRNA had moderate viral loads (1.5×10^5^–8.5×10^5^ PFU/tissue). At the same time point, we also observed undetectable or low viral loads (1.1×10^2^–3.4×10^2^ PFU/tissue) in the lungs of most groups immunized with mRNA encoding SARS-CoV-2 immunogens, the exception being mice immunized with 1 µg of mRNA encoding secreted Rpk9-I53-50A (1.5×10^4^ PFU/tissue) (**Fig. 5f**). As expected, mice immunized with PBS and luciferase mRNA had high viral loads (3.4×10^7^–4.1×10^7^ PFU/tissue) two days post infection. Four days post infection, we observed a decrease in lung viral titers across all groups. All groups immunized with mRNA encoding SARS-CoV-2 immunogens had undetectable viral loads, whereas mice immunized with PBS and luciferase mRNA maintained moderate viral loads (1.5×10^5^–4.0×10^5^ PFU/tissue). Altogether, these results indicate that mRNA-launched RBD nanoparticles elicit more protective antibodies and provide better protection against vaccine-mismatched challenge than COVID-19 mRNA vaccines encoding membrane-anchored S-2P or a secreted RBD trimer.

## Discussion

The COVID-19 pandemic established that mRNA vaccines can be safe, effective, and rapidly scalable. This success was made possible by decades of research focused on optimizing both the mRNA backbone (i.e., nucleoside modifications^64–66^, 5′ capping strategies^67–69^, UTRs^70–72^, etc.) and the delivery vehicle (i.e., LNP formulations^73–75^). However, there is another aspect of mRNA vaccine development that has been surprisingly understudied: how to best design the encoded protein immunogens to improve their immunogenicity while ensuring they remain compatible with mRNA delivery. Licensure of COVID-19 mRNA vaccines encoding engineered prefusion-stabilized SARS-CoV-2 Spike protein^19,21^ provides a strong rationale for protein design as a major consideration in mRNA vaccine development.

Here, we applied protein design to develop an RBD nanoparticle immunogen compatible with delivery as an mRNA vaccine. Specifically, we used a one-component (i.e., homomeric) protein nanoparticle previously designed for optimal secretion^14^ to display a SARS-CoV-2 RBD antigen designed for improved stability and expression^15^. When produced in cell culture, the resultant RBD nanoparticle was found to efficiently secrete and self-assemble while maintaining its structure and antigenicity. When delivered as an mRNA vaccine, the RBD nanoparticle elicited neutralizing antibody responses with improved potency, breadth, and protective capacity compared to mRNA-delivered membrane-anchored Spike trimers and secreted RBD trimers. Additionally, the RBD nanoparticle induced robust antigen-specific CD8 T cell responses when delivered as an mRNA vaccine, but not as an adjuvanted protein vaccine. These results highlight how computationally designed mRNA-launched protein nanoparticle vaccines retain the advantageous features of both protein nanoparticle vaccines (i.e., enhanced B cell activation and expansion^7,76^, superior trafficking to lymph nodes and B cell follicles^77–79^, and improved antigen stability^80,81^) and mRNA-LNP vaccines (i.e., intracellular processing and MHC I presentation of antigen^82^, inherent adjuvanticity of LNPs^13,83–86^).

Previously, genetically delivered protein nanoparticle vaccines have been based on naturally occurring scaffolds such as ferritin and lumazine synthase^34–37,87–90^. Here, we used I3-01NS, a protein nanoparticle designed and optimized for secretion with Rosetta^14,91,92^, to demonstrate proof-of-concept for computationally designed mRNA-launched protein nanoparticle vaccines. We envision two paths forward for this technology, each with its respective advantages. On the one hand, robust and versatile protein nanoparticles like I3-01NS, ferritin, and lumazine synthase are broadly applicable “platforms” that could be used to generate a number of vaccines. For example, I3-01 has previously been used to generate protein nanoparticle vaccine candidates displaying several different antigens via genetic fusion^93–97^. Our work suggests that I3-01NS may prove similarly versatile as a platform for mRNA-launched nanoparticle vaccines. However, we note that the initial failure of RBD-I3-01NS to secrete indicates that further work will be required to fully understand and optimize the relationship between displayed antigens, the I3-01NS scaffold, and secretion. On the other hand, the structural features of I3-01NS and the small number of other naturally occurring proteins that have been used as mRNA-launched nanoparticle vaccine scaffolds are relatively unalterable. This is a considerable limitation, as several recent studies have shown that the detailed geometry of antigen presentation can significantly impact B cell activation and vaccine-elicited antibody responses^7,98–100^. Computational methods that allow the design of novel self-assembling proteins with atomic-level accuracy overcome this limitation by providing a route to genetically delivered nanoparticle immunogens with custom structural and functional features^101^. The recent development of powerful machine learning-based methods for protein structure prediction and design^102–104^ will facilitate these efforts and may enable the development of nanoparticle vaccine scaffolds that are truly tailored to specific antigens.

In conclusion, our work demonstrates the utility of, and lays the foundation for, computationally designed mRNA-launched protein nanoparticle vaccines. We anticipate that this technology will be useful in designing vaccines against various viral, bacterial, and parasitic pathogens, and may be particularly valuable for pandemic prevention, preparedness, and response^105^.

## Supporting information

Hendricks_et_al_statistical_analyses

Hendricks_et_al_Source_Data_Main_Figs

Hendricks_et_al_Source_Data_Extended_Data_Figs

## Acknowledgements

The authors gratefully acknowledge Lynda Stuart, Holger Kanzler, and Harry Kleanthous for helpful discussions; Andrew Borst, Rebecca Skotheim, Annika Philomin, Kenneth Carr, and Connor Weidle for maintaining the electron microscopy facilities at the Institute for Protein Design; Michael Murphy, Claire Sydeman, Maggie Ahlrichs, Craig Dobbins, and Alexis Hand for maintaining and providing mammalian cells; Rashmi Ravichandran, Samuel Wrenn, Elizabeth Kepl, Brooke Fiala, and Natalie Brunette for assistance with protein production; Lauren Carter, Kandise VanWormer, Ratika Krishnamurty, Kristina Herrera, Madison Kennedy, and Lance Stewart for laboratory and administrative support; and Luki Goldschmidt and Patrick Vecchiato for building and maintaining the computing infrastructure at the Institute for Protein Design. This work was funded by the Bill & Melinda Gates Foundation (INV-010680, INV-043758, INV-018675, and INV-003875), the National Institute of Allergy and Infectious Disease (1P01AI167966, DP1AI158186, 75N93022C00036, R01AI153064, P01AI158571, and P01AI172531), an Investigators in the Pathogenesis of Infectious Disease Awards from the Burroughs Wellcome Fund, the National Institutes of Health (S10OD032290), and the Audacious Project at the Institute for Protein Design. D.V. is an Investigator of the Howard Hughes Medical Institute and the Hans Neurath Endowed Chair in Biochemistry at the University of Washington. G.G.H. was supported by a generous gift from the Higgins family.

## Author contributions

G.G.H., D.E., D.V., A.S., and N.P.K. designed the study. A.C.W., D.E., D.V., and N.P.K. designed immunogens. G.G.H. and D.E. produced and characterized immunogens. G.G.H., J.Y.J.W., M.C.M., and A.V. produced and provided proteins. G.G.H., S. Chan, and C.M. performed transfections. C.W.C. performed electron microscopy experiments. G.G.H., M.J.N., M.M., A.C.W., J.L., and S. Cheng performed serology experiments. C.T., M.R.J., and E.M.L. conducted the protein vs mRNA immunogenicity study. L.G., H.H., S.Y.W., and V.L. conducted the T cell study and performed intracellular cytokine staining experiments. N.J.C., M.L.H., J.M.P., and A.S. conducted the challenge studies and performed plaque assays. R.S.B., B.P., and A.S. provided oversight of animal studies. H.M. and N.P. produced and provided luciferase mRNA. P.J.C.L., M.M.H.S., and Y.K.T. performed LNP encapsulation experiments. G.G.H., L.G., M.J.N., N.J.C., M.L.H., J.M.P., M.M., A.C.W., C.W.C., H.H., S.Y.W., V.L., R.S.B., B.P., D.V., A.S., and N.P.K. analyzed the data. G.G.H. and N.P.K. prepared figures and wrote the manuscript with input from all authors.

## Methods

### Plasmid construction

Wild-type RBD and stabilized Rpk9 sequences were genetically fused to the N termini of the I3-01NS nanoparticle component using linkers of 16 glycine and serine residues. The sequences also contained an N-terminal signal peptide (MGILPSPGMPALLSLVSLLSVLLMGCVAETGT). The sequences were codon-optimized for human cell expression and cloned into pCMV/R using the XbaI and AvrII restriction sites by Genscript. The plasmids for hACE2-Fc, CR3022, Rpk9-I53-50A, S-2P-foldon, HexaPro, I3-01NS, I53-50A, I53-50B, and CV30 were produced as previously described^14,15,28,52,111^. Plasmids were transformed into the NEB 5α strain of *E. coli* (New England Biolabs) for subsequent DNA extraction (QIAGEN Plasmid PlusTM Maxi Kit protocol) to obtain plasmid for transient transfection into Expi293F cells. The amino acid sequences of all proteins used for immunizations can be found in **Supplementary Table 1**.

### Protein production and purification

Expi293F cells were grown in suspension and passaged according to manufacturer protocols (ThermoFisher). Cells at 3.0×10^6^ cells/mL were transfected with 1 μg of purified plasmid DNA mixed with 3 μg of PEI-MAX per mL cell culture. Supernatant was harvested 72 h post-transfection by centrifugation for 5 minutes at 4,100 *g* followed by the addition of PDADMAC solution (Sigma Aldrich), a second centrifugation for 5 minutes at 4,100 *g*, and sterile filtration.

To purify Rpk9-I3-01NS for animal studies, the supernatant was adjusted to 50 mM Tris (pH 8.0). Galanthus Nivalis Gel (GNA) immobilized lectin conjugated resin (EY Laboratories) was rinsed into magnesium- and calcium-free (D)PBS (Gibco) using a gravity column and then added to the supernatant, followed by overnight shaking at 4°C. The resin was collected using a gravity column and washed with Lectin Wash Buffer (50 mM Tris (pH 8.0), 150 mM NaCl, 100 mM Arginine (pH 8.0), 5% (v/v) glycerol, and 0.02% (w/v) NaN_3_). Protein was eluted using Lectin Elution Buffer (50 mM Tris (pH 8.0) 150 mM NaCl, 100 mM Arginine (pH 8.0), 5% (v/v) glycerol, 0.02% (w/v) NaN_3_, and 1 M Methyl-α-D-mannopyranoside). Eluates were concentrated and applied to a Superose 6 Increase 10/300 GL column pre-equilibrated with Sizing Buffer (50 mM Tris (pH 8.0), 150 mM NaCl, 100 mM Arginine (pH 8.0), and 5% (v/v) glycerol) for preparative SEC. Peaks corresponding to RBD nanoparticles were identified based on elution volume, pooled together, and sterile-filtered. Protein samples were stored at 4°C or -80°C until use.

When purifying Rpk9-I3-01NS for supernatant ELISA standard curves, the 5% (v/v) glycerol and 0.02% (w/v) NaN_3_ were omitted from the Lectin Wash Buffer and Lectin Elution Buffer. Purification and production of hACE2-Fc, CR3022, Rpk9-I53-50A, S-2P-foldon, HexaPro, I3-01NS, I53-50A, I53-50B, and CV30 was performed as previously described^14,15,28,52,111^.

### Ultraviolet-visible spectrophotometry (UV/vis)

Protein samples were applied to a 10 mm, 50 μL quartz cell (Starna Cells, Inc.) and absorbance was measured from 180 to 1000 nm using an Agilent Cary 8454 spectrophotometer. Net absorbance at 280 nm, obtained from measurement and single reference wavelength baseline subtraction, was used with calculated extinction coefficients and molecular weights to obtain protein concentration. Samples were diluted with respective blanking buffers to obtain an absorbance between 0.1 and 1.0. All data produced from the UV/vis instrument was processed in the 845x UV/visible System software.

### Dynamic light scattering (DLS)

Hydrodynamic diameter (D_h_) and polydispersity (Pd) was measured on an UNcle Nano-DSF (UNchained Laboratories) at 25°C. Sample was applied to a 8.8 μL quartz capillary cassette (UNi, UNchained Laboratories) and measured with 10 acquisitions of 5 s each, using auto-attenuation of the laser. Protein concentration (ranging from 0.1–1.0 mg/mL) and increased viscosity due to the inclusion of 5% v/v glycerol in the buffer was accounted for by the UNcle Client software.

### Negative stain electron microscopy (nsEM)

Protein samples (3–6 μL, 0.05–0.1 mg/mL) were applied to glow-discharged 300-mesh copper grids (Ted Pella) and stained with uranyl formate (0.75–2% (w/v)). Data were collected using a 120 kV Talos L120C transmission electron microscope (Thermo Scientific) with a BM-Ceta camera. CryoSPARC^112^ was used for CTF correction, particle picking and extraction, and 2D classification.

### Biolayer interferometry (BLI)

Binding of hACE2-Fc, CR3022, and S309 (Abcam; ab289796) to RBD nanoparticles was analyzed using an Octet Red 96 System (Pall FortéBio/Sartorius) at ambient temperature with shaking at 1000 rpm. Protein samples were diluted to 100 nM in Sartorius Octet Kinetics Buffer 10× (diluted to 1×). Kinetics Buffer, antibody, receptor, and immunogen were then applied to a black 96-well Greiner Bio-one microplate at 200 μL per well. Protein A biosensors were first hydrated for 10 min, then equilibrated in Kinetics Buffer for 30 s. The tips were then dipped into either hACE2-Fc, CR3022, and S309 diluted to 10 μg/mL in Kinetics Buffer for 300 s, then transferred back into Kinetics Buffer for 60 s to reach a baseline. The association step was performed by dipping the loaded biosensors into the immunogens for 300 s, and the subsequent dissociation step was performed by dipping the biosensors back into the Kinetics Buffer used to baseline for an additional 300 s. The data were baseline subtracted prior to plotting using the FortéBio analysis software.

### Supernatant enzyme-linked immunosorbent assays (ELISA)

Supernatants were serially diluted 1:2 in Expi293 Expression Medium (Gibco). Purified protein (Rpk9-I3-01NS for RBD nanoparticles; Rpk9-I53-50A for RBD trimers) was diluted in Expi293 Expression Medium (Gibco) to create a standard curve at known concentrations. For each immunogen, 100 μL of the cognate supernatant and standard curve dilutions were plated onto 96-well Nunc Maxisorp (ThermoFisher) plates. Plates were incubated at 25°C for 1 h then washed 3× using a plate washer (BioTek). Plates were blocked with 200 μL of 5% non-fat milk in TBST (25 mM Tris (pH 8.0), 150 mM NaCl, 0.05% (v/v) Tween-20) for 1 h at 25°C. Plates were washed 3× in TBST, then 100 μL of 4 μg/mL CV30^113^, a RBD-directed mAb, was added to each well and incubated at 25°C for 1 h. Plates were washed 3× in TBST, then anti-human (Southern Biotech) horseradish peroxidase-conjugated antibodies were diluted 1:5,000 and 100 μL was added to each well and incubated at 25°C for 30 min. Plates were washed 3× in TBST and 100 μL of TMB (SeraCare) was added to every well for 2 min at room temperature. The reaction was quenched with the addition of 100 μL of 1 N HCl. Plates were immediately read at 450 nm on an Epoch2 plate reader. The data was plotted and fit in Prism (GraphPad) using nonlinear regression sigmoidal, 4PL, X is concentration to determine the concentration of secreted protein.

### Endotoxin measurements

Endotoxin levels in protein samples were measured using the EndoSafe PTS System (Charles River). Samples were diluted 1:50 or 1:100 in Endotoxin-free LAL reagent water, and applied into wells of an EndoSafe LAL reagent cartridge. Endotoxin values were reported as EU/mL with the dilution factor automatically back-calculated. Our threshold for samples suitable for immunization was <5 EU/dose. Protein samples with endotoxin levels above this threshold were spiked with 0.75% w/v 3-[(3-cholamidopropyl)dimethylammonio]-1-propanesulfonate (CHAPS) and dialyzed three times against respective Sizing Buffers in a hydrated 20K molecular weight cutoff dialysis cassette (ThermoFisher).

### mRNA design and *in vitro* transcription

Membrane-anchored S-2P mRNA was designed to be identical to the publicly available, reverse engineered nucleic acid sequence of Pfizer/BioNTech’s BNT162b2 vaccine^114,115^. Rpk9-I53-50A and Rpk9-I3-01NS mRNAs were designed by retaining the UTRs and polyA tail from the Pfizer/BioNTech BNT162b2 vaccine but replacing the Spike-encoding open-reading frame (ORF) for the codon-optimized ORF of the Rpk9-I53-50A and Rpk9-I3-01NS pCMV/R expression plasmids, respectively. *In vitro* transcription of membrane-anchored S-2P, Rpk9-I3-01NS, and Rpk9-I53-50A mRNA was performed by TriLink Biotechnologies using previously described standard protocols^69,116^, N1-methylpseudouridine-5’-triphosphate (m1Ψ-5′-triphosphate) (TriLink #N-1081) instead of uridine-5’-triphosphate (UTP), and the CleanCap Reagent AG (TriLink #N-7113) for co-transcriptional capping.

The firefly luciferase (Luc)-encoding mRNA was produced as described^117^. Briefly, the Luc sequence was codon-optimized, synthesized, and cloned into an mRNA production plasmid. The mRNA was transcribed to contain a 101 nucleotide-long poly(A) tail. N1-methylpseudouridine-5’-triphosphate (m1Ψ-5′-triphosphate) (TriLink #N-1081) instead of uridine-5’-triphosphate (UTP) was used to generate modified nucleoside-containing mRNA. Capping of the in vitro transcribed mRNAs was performed co-transcriptionally using the trinucleotide cap1 analog, CleanCap (TriLink #N-7413). mRNA was purified by cellulose (Sigma-Aldrich #11363-250G) purification. Luciferase mRNAs were analyzed by agarose gel electrophoresis and were stored frozen at −20 °C.

Nucleic acid sequences for the ORFs of all mRNA constructs used for immunizations can be found in Supplementary Table 2.

### Lipid nanoparticle (LNP) encapsulation of mRNAs

Lipid nanoparticles used in this study were similar in composition to those previously described^118,119^ and contain ionizable lipids proprietary to Acuitas Therapeutics (pKa ranging from 6.0–6.5). Nucleoside-modified mRNAs were encapsulated in LNPs using a self-assembly process in which an aqueous solution of mRNA (pH 4.0) was rapidly mixed with an aqueous solution of lipids (ionizable lipid/distearoylphosphatidylcholine (DSPC)/cholesterol/PEGylated lipid) dissolved in ethanol. The resultant mRNA-LNPs were characterized at Acuitas Therapeutics for their size, polydispersity, and encapsulation efficiency, then stored at -80°C until use.

### Formulation of proteins and mRNA-LNPs for immunizations

Prior to all immunizations, protein immunogen suspensions were diluted in Sizing Buffer and gently mixed 1:1 (v/v) with AddaVax adjuvant (Invivogen) to reach a final dose of 0.9 μg of RBD per mouse for Rpk9-based constructs or 5 μg of Spike per mouse for S-2P-foldon, which comprise equimolar amounts of RBD antigen. Prior to all immunizations, mRNA-LNP suspensions were diluted with magnesium- and calcium-free Dulbecco’s Phosphate Buffered Saline (Gibco) to reach a final dose of 0.2, 1, or 5 μg mRNA per mouse.

### Protein and mRNA-LNP immunogenicity study

Female BALB/c mice were purchased from Envigo (order code 047) at 7 weeks of age and were maintained in a specific pathogen-free facility within the Department of Comparative Medicine at the University of Washington, Seattle, accredited by the Association for Assessment and Accreditation of Laboratory Animal Care (AAALAC). Animal experiments were conducted in accordance with the University of Washington’s Institutional Animal Care and Use Committee under protocol 4470-01. Mice of 8 weeks of age were injected intramuscularly into the quadriceps muscle of both hind legs with 50 μL per injection site under isoflurane anesthesia. Blood was collected via submental venous puncture and rested in serum separator tubes (BD # 365967) at room temperature for 30 min to allow for coagulation. Serum was separated from hematocrit via centrifugation at 2,000 *g* for 10 min. Complement factors and pathogens in isolated serum were heat-inactivated via incubation at 56°C for 60 min. Serum was stored at 4°C or -80°C until use.

### Serum enzyme-linked immunosorbent assays (ELISA)

For antigen-specific ELISAs, 100 μL of 2 μg/mL HexaPro was plated onto 96-well Nunc Maxisorp (ThermoFisher) plates in 50 mM Tris (pH 8.0), 150 mM NaCl, 0.25% (v/v) L-Histidine, 5% (v/v) glycerol. For scaffold-specific ELISAs, 100 μL of 2 μg/mL I3-01NS, I53-50A, or I53-50 was plated onto 96-well Nunc Maxisorp (ThermoFisher) plates in TBS (25 mM Tris (pH 8.0), 150 mM NaCl). Plates were incubated at 25°C for 1 h then washed 3× in TBST using a plate washer (BioTek). Plates were blocked with 200 μL of 5% non-fat milk in TBST for 1 h at 25°C. Plates were washed 3× in TBST and 1:3 or 1:5 serial dilutions of mouse sera were made in 5% non-fat milk in TBST starting at 1:50 and incubated at 25°C for 1 h. Plates were washed 3× in TBST, then anti-mouse (Cell Signaling Technologies) horseradish peroxidase-conjugated antibodies were diluted 1:2,000 and 100 μL was added to each well and incubated at 25°C for 30 min. Plates were washed 3× in TBST and 100 μL of TMB (SeraCare) was added to every well for 3 min at room temperature. The reaction was quenched with the addition of 100 μL of 1 N HCl. Plates were immediately read at 450 nm on an Epoch2 plate reader. The data was plotted and fit in Prism (GraphPad) using nonlinear regression sigmoidal, 4PL, X is log(concentration) to determine the reciprocal ED_50_ values.

### Pseudovirus production

The full-length D614G Wuhan-Hu-1 and Omicron BA.2 S constructs with a 21-amino-acid C-terminal deletion used for pseudovirus assays were previously described^120–122^. The full-length Omicron BA.2 construct containing a 21-amino-acid C-terminal deletion was codon optimized, synthesized, and inserted the HDM vector by Genscript. Pseudotyped VSV was produced as previously described^120–122^. In brief, HEK293T cells were split into poly-D-lysine-coated 15-cm plates and grown overnight until they reached approximately 70–80% confluency. The cells were washed 3 times with Opti-MEM (Gibco) and transfected with either the D614G Wuhan-Hu-1 or Omicron BA.2 S constructs using Lipofectamine 2000 (Life Technologies). After 4–6 h, the medium was supplemented with an equal volume of DMEM supplemented with 20% FBS and 2% penicillin-streptomycin. The cells were incubated for 20–24 h, washed 3 times with DMEM, and infected with VSVΔG-luc. Two hours after VSVΔG-luc infection, the cells were washed an additional five times with DMEM. The cells were grown in DMEM supplemented with anti-VSV-G antibody (I1-mouse hybridoma supernatant diluted 1:25, from CRL-2700, ATCC) for 18–24 h, after which the supernatant was harvested and clarified by low-speed centrifugation at 2,500g for 10 min. The supernatant was then filtered (0.45 μm) and virus stocks were concentrated 10 times using a 30 kDa centrifugal concentrator (Amicon Ultra). The pseudotyped viruses were then aliquoted and frozen at −80 °C.

### Pseudovirus neutralization assays

VeroE6-TMPRSS2^123^ were split into white-walled, clear-bottom 96-well plates (Corning) and grown overnight until they reached approximately 70% confluency. Plasma was diluted in DMEM and serially diluted in DMEM at a 1:3 dilution thereafter. Pseudotyped VSV was diluted at a 1:20 to 1:100 ratio in DMEM and an equal volume was added to the diluted plasma. The virus–plasma mixture was incubated for 30 min at room temperature and added to the Vero E6-TMPRSS2 cells. After two hours, an equal volume of DMEM supplemented with 20% FBS and 2% penicillin-streptomycin was added to the cells. After 20–24 h, ONE-Glo EX (Promega) was added to each well and the cells were incubated for 5 min at 37 °C. Luminescence values were measured using a BioTek Synergy Neo2 plate reader. Luminescence readings from the neutralization assays were normalized and analyzed using GraphPad Prism 9.4.1. The Relative Light Unit (RLU) values recorded from uninfected cells were used to define 100% neutralization and RLU values recorded from cells infected with pseudovirus without plasma were used to define 0% neutralization. Reciprocal ID_50_ values were determined in Prism (GraphPad) from the normalized data points using a log(inhibitor) versus normalized response–variable slope model to generate the curve fits. Each neutralization experiment was conducted at least twice using independently produced batches of pseudoviruses. Representative data from a single experiment are provided.

### Evaluation of T cell responses

Eight- to twelve-week-old C57BL/6 mice were immunized intramuscularly at weeks 0 and 3 with diluted adjuvanted protein immunogen or diluted mRNA-LNP. Lungs and spleens from immunized animals were harvested at week 6 and lymphocyte isolation from both tissues was performed. Lymphocytes from lungs and spleens were plated in a 96-well round-bottom plate at a density of 1 × 10^6^ cells/mL, in 200 μL final volume with complete T cell media (containing RPMI-1640, FBS, Penicillin/Streptomycin, non-essential amino acids, β-mercaptoethanol, HEPES). Cells were stimulated with an overlapping peptide pool derived from SARS-CoV-2 Spike protein (Genscript). Following 2 hours of culture, Brefeldin A was added, and cells were left in culture for 8 more hours. Following the stimulation, cells were stained with an extracellular antibody cocktail containing anti-CD3 (BV785; BioLegend #100355, 1:50 dilution), anti-CD4 (BV650; BioLegend #100555; 1:200 dilution), anti-CD8 (BV711; BD #563046; 1:200 dilution), anti-CD45 (BUV395, BD # 564279). Cells were fixed and permeabilized with BD Cytofix/Cytoperm, then stained intracellularly with a cocktail of antibodies containing anti-IFNγ (APC; BioLegend #505810; 1:100 dilution), anti-TNFα (FITC; BioLegend #506304, 1:100 dilution), anti-IL-2 (PE; BioLegend #503808, 1:100 dilution), and anti-IL-4 (PerCPCy5.5; BioLegend #504124, 1:100 dilution). Cells were analyzed with an LSRII.UV analyzer at the Stanford Shared FACS Facility (SSFF).

### Viruses

The mouse-adapted viruses (Wuhan-Hu-1 SARS-CoV-2 MA10 and Omicron BA.5 SARS-CoV-2) and authentic viruses (D614G Wuhan-Hu-1 SARS-CoV-2 and Omicron BA.5 SARS-CoV-2) nanoluciferase recombinant strains used in this study were described previously^56,63,124,125^. All virus stocks were propagated in VeroE6-TMPRSS2 cells and subjected to next-generation sequencing (Illumina) to confirm the introduction and stability of substitutions.

### Wuhan-Hu-1 SARS-CoV-2 MA10 challenge

Animal studies were carried out in accordance with the Institutional Animal Care and Use Committee at UNC Chapel Hill (protocol number 23-085) and were performed in approved biosafety level 3 (BSL-3) facilities. Ten-week-old female BALB/c mice (Inotiv, code 047; groups of 4-6 mice per group per time point) were vaccinated intramuscularly into the quadriceps muscle of both hind legs with 25 μL of diluted mRNA-LNP or PBS per injection site. Five weeks after vaccination, serum was collected and mice were moved into BSL-3 for homologous challenge on week 6. Briefly, mice were anesthetized with a mixture of ketamine/xylazine, then inoculated intranasally with 1×10^5^ PFU of Wuhan-Hu-1 SARS-CoV-2 MA10. Mice were then monitored daily for clinical signs of disease, weight loss, and mortality. At 2 and 4 days post infection (dpi), mice were euthanized via isoflurane overdose, and the congestion score was assessed (for 4 dpi only). The inferior lung lobe and nasal turbinates (for 2 dpi only) were collected in PBS with glass beads and stored at -80°C for viral titer determination via plaque assay, as previously described^56^. Briefly, harvested tissues were homogenized in PBS and serial dilutions of the clarified homogenates were added to a confluent monolayer of Vero E6 cells followed by agarose overlay. Plaques were then visualized via staining with Neutral Red dye and counted on 3 dpi.

### Omicron BA.5 SARS-CoV-2 challenge

Animal studies were carried out in accordance with the Institutional Animal Care and Use Committee at UNC-Chapel Hill (protocol number 23-085) and were performed in approved biosafety level 3 (BSL-3) facilities. On weeks 0 and 4, ten-week-old female BALB/c mice (Inotiv, code 047; groups of 4-5 mice per group per time point) were vaccinated intramuscularly into the quadriceps muscle of both hind legs with 25 μL of diluted mRNA-LNP or PBS per injection site. Eight weeks after vaccination, serum was collected and mice were moved into BSL-3 for heterologous challenge on week 9. Briefly, mice were anesthetized with a mixture of ketamine/xylazine, then inoculated intranasally with 1×10^5^ PFU of Omicron BA.5 SARS-CoV-2. Mice were then monitored daily for clinical signs of disease, weight loss, and mortality. At 2 and 4 days post infection (dpi), mice were euthanized via isoflurane overdose and the congestion score was assessed. The inferior lung lobe and nasal turbinates (for 2 dpi only) were collected in PBS with glass beads and stored at -80°C for viral titer determination via plaque assay, as previously described^56^. Briefly, harvested tissues were homogenized in PBS and serial dilutions of the clarified homogenates were added to a confluent monolayer of Vero E6 cells followed by agarose overlay. Plaques were then visualized via staining with Neutral Red dye and counted on 3 dpi.

### Authentic virus neutralization assays

Pre-challenge sera were serially diluted (1:20 initially, followed by a 5-fold dilution) in 96-well round-bottom plates (Corning 3799) and incubated at a 1:1 ratio with D614G Wuhan-Hu-1 SARS-CoV-2 or Omicron BA.5 SARS-CoV-2 expressing nanoluciferase for 1 hour at 37°C, 5%CO2. Following incubation, the serum-virus complexes were added, in duplicate, to confluent monolayers of Vero C1008 cells (ATCC, cat# CRL-1586) and incubated for 26-30 hours, depending on the virus. After incubation, RLU values were measured with the Nano-Glo Luciferase Assay System (Promega) according to the manufacturer’s protocol. Neutralization titers were calculated as the dilution at which a 50% reduction in RLU values was observed relative to the virus-only control. Curve fits were generated using a custom macro constrained at the top (100% neutralization) but not the bottom (0% neutralization). Assays were performed once with technical duplicates.

### Statistical analysis

All statistical analyses are available as **Supplementary Information**. Serological data, intracellular cytokine staining data, and nasal turbinate viral titer data were analyzed using one-way ANOVA followed by Tukey’s multiple comparisons. Congestion scores were analyzed using one-way (for Fig. 4d) or two-way (for Fig. 5d) ANOVA followed by Tukey’s multiple comparisons. Weight loss data was analyzed using mixed effect analysis followed by Tukey’s multiple comparisons. Lung viral titer data was analyzed using two-way ANOVA followed by Tukey’s multiple comparisons. When comparing only two groups, statistical significance was calculated using a two-tailed non-parametric Mann-Whitney test.

## Data Availability

All images and data were generated and analyzed by the authors. **Source Data** for Figs.1b, c, e; 2b-d; 3b, c; 4b-f; 5b-f and Extended Data Figs.1a-h; 5b; 7b, c; 8b are provided with this paper. Other data will be made available by the corresponding author upon reasonable request.

## Extended Data Figures

**Extended Data Fig. 1.**
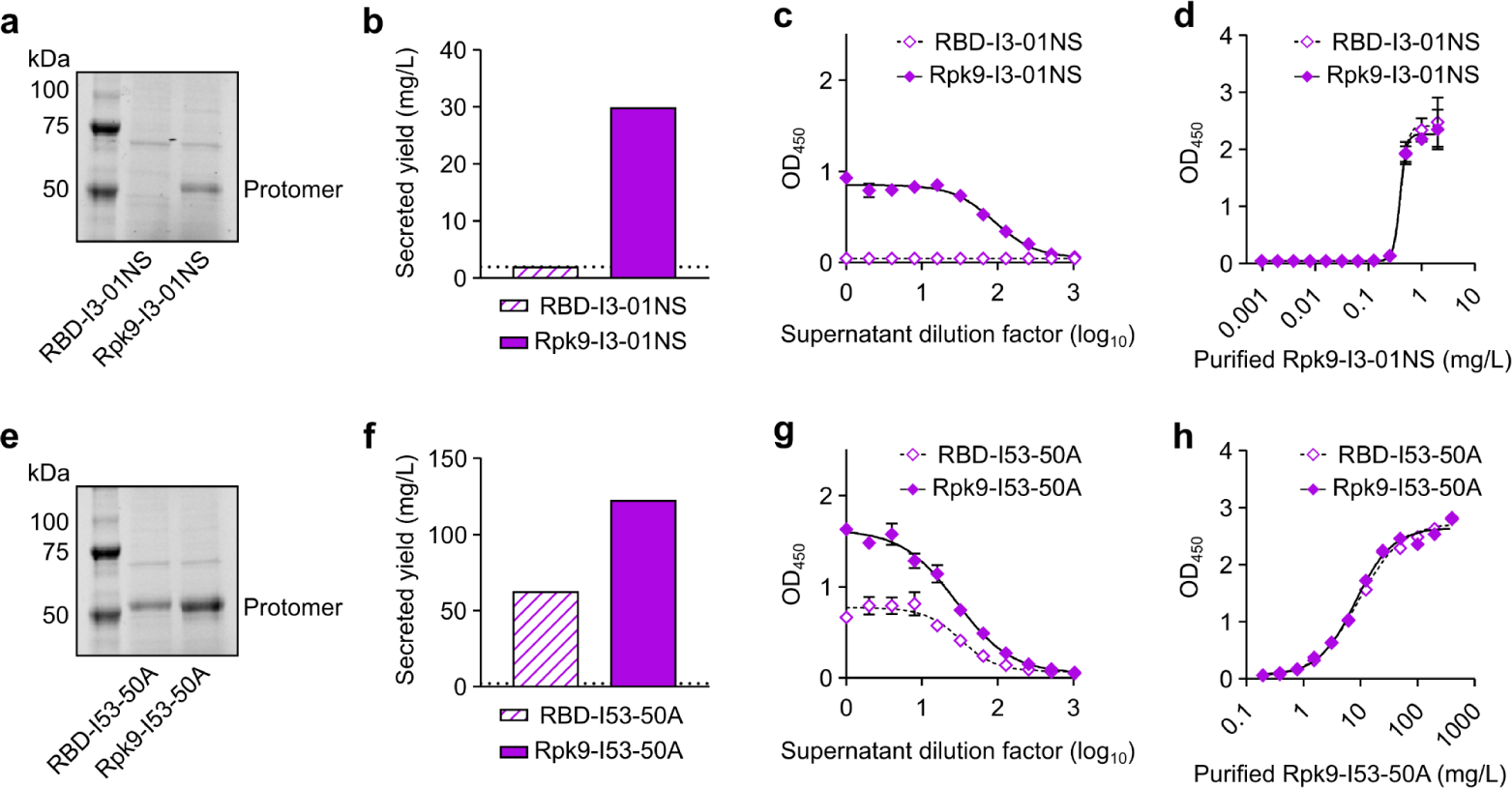
Quantification of secreted RBD nanoparticles and RBD trimers. **a,** Reducing SDS-PAGE of supernatants from Expi293F cells after expression of RBD nanoparticles. **b,** Secreted yield of RBD nanoparticles, as determined by cell supernatant ELISAs. **c,** RBD nanoparticle supernatant binding against CV30. **d,** Purified Rpk9-I3-01NS binding against CV30 was used to quantify the secreted yield of both supernatants. The standard curves generated for analysis are shown for each supernatant. **e,** Reducing SDS-PAGE of supernatants from Expi293F cells after expression of RBD trimers. **f,** Secreted yield of RBD trimers, as determined by supernatant ELISAs. **g,** RBD trimer supernatant binding against CV30. **h,** Purified Rpk9-I53-50A binding against CV30 was used to quantify the secreted yield of both supernatants. The standard curves generated for analysis are shown for each supernatant. Note the very different x axis scales in panels **d** and **h**, highlighting that nanoparticles and trimers require distinct standard curves for accurate quantitation. **a-h,** Representative data are shown from one of three biological replicates for each construct. **b, f** The dotted horizontal line represents the limit of detection for the assay. **c-d, g-h** Error bars represent mean ± SD of two technical replicates.

**Extended Data Fig. 2.**
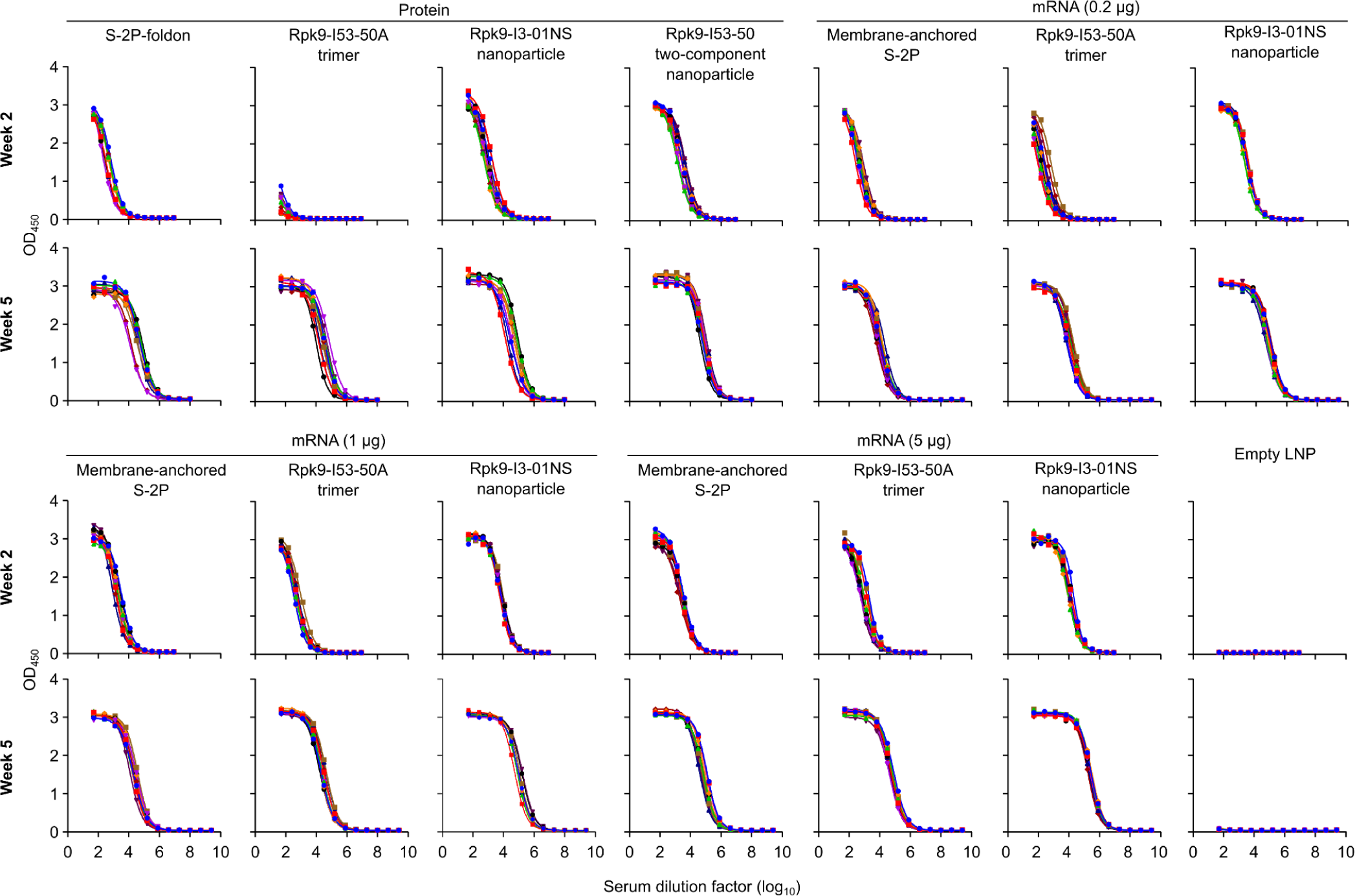
Raw ELISA data and fits used to determine titers shown in Figure 2. Serum binding against Wuhan-Hu-1 SARS-CoV-2 S HexaPro. Each color represents an individual animal.

**Extended Data Fig. 3.**
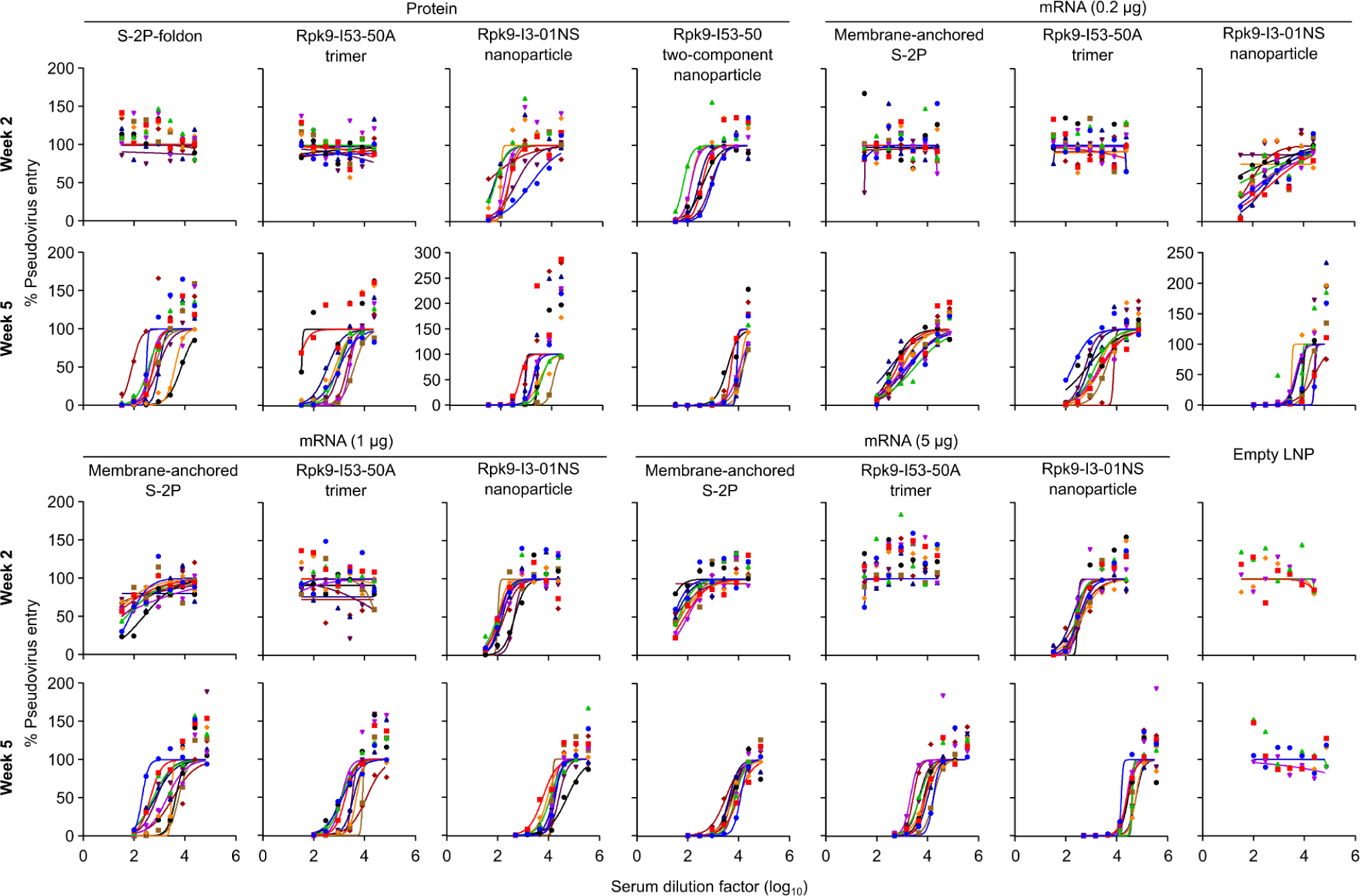
Dose-response curves of D614G Wuhan-Hu-1 SARS-CoV-2 pseudovirus neutralization, related to Figure 2. Serum neutralizing activity against VSV pseudotyped with D614G Wuhan-Hu-1 SARS-CoV-2 S. Each color represents an individual animal. Some groups do not have data for all 10 mice due to a lack of available sera (for week 2: n=9 for S-2P-foldon, n=8 for Rpk9-I53-50, n=9 for Rpk9-I3-01NS (protein), and n=4 for Empty LNP). The y axis range and scale of each plot correspond to the leftmost graph in the row unless otherwise indicated.

**Extended Data Fig. 4.**
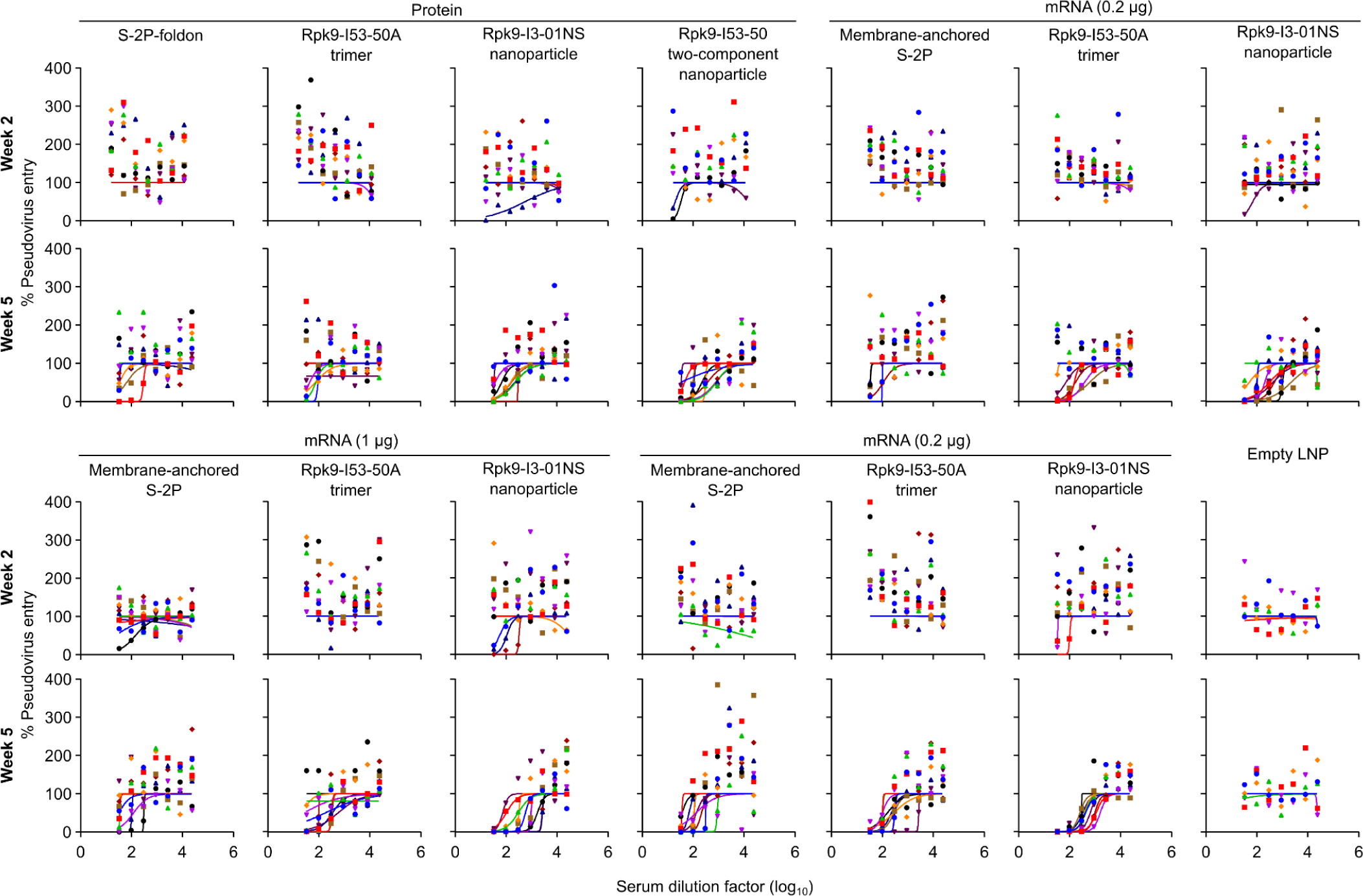
Dose-response curves of Omicron BA.2 SARS-CoV-2 pseudovirus neutralization, related to Figure 2. Serum neutralizing activity against VSV pseudotyped with Omicron BA.2 SARS-CoV-2 S. Each color represents an individual animal. Some groups do not have data for all 10 mice due to a lack of available sera (for week 2: n=9 for S-2P-foldon, n=8 for Rpk9-I53-50, n=9 for Rpk9-I3-01NS (protein)).

**Extended Data Fig. 5.**
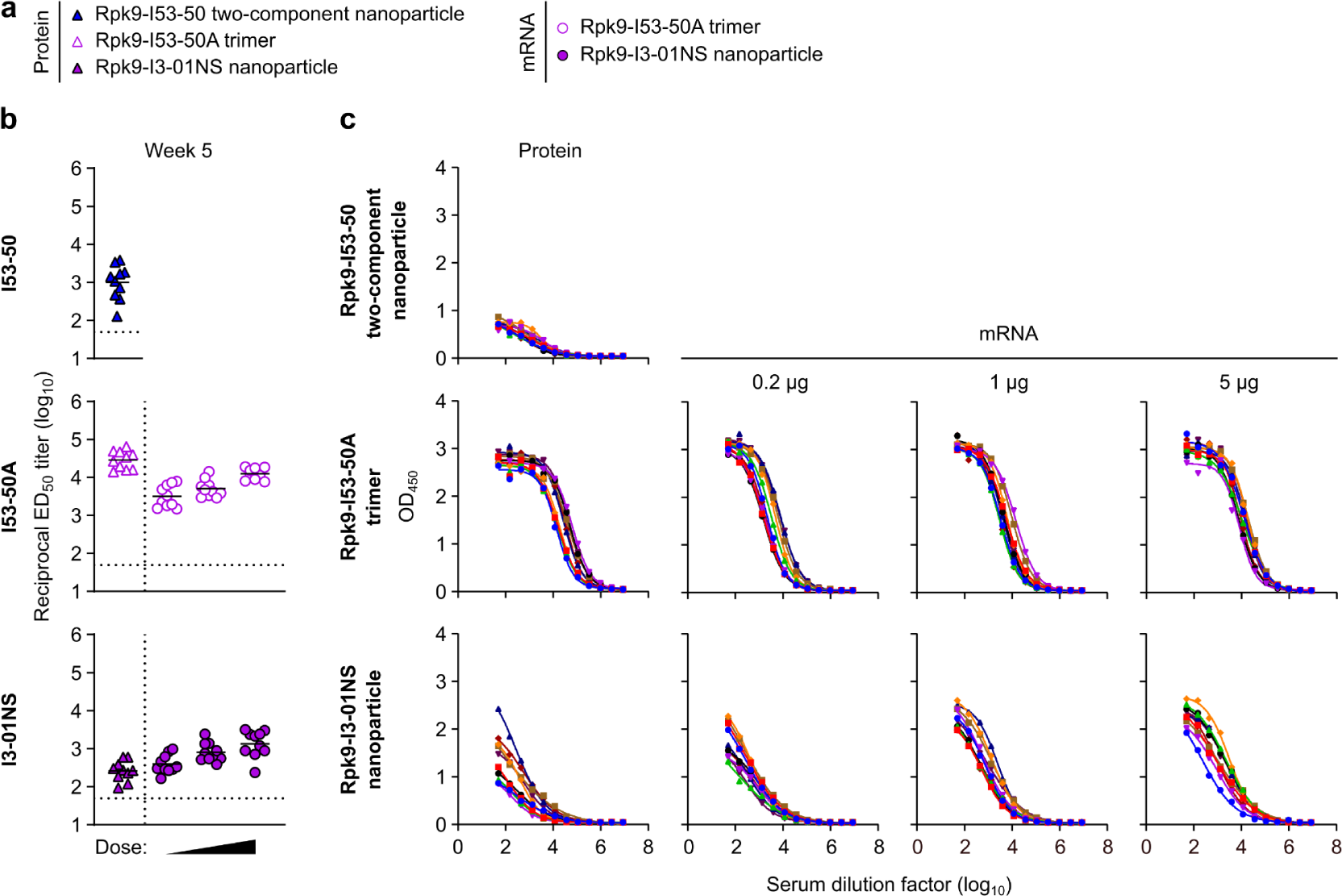
Anti-scaffold antibody titers in BALB/c mice two weeks post-boost. **a,** Groups assessed for anti-scaffold responses at week 5 (from mice in Fig. 2). **b,** Serum antibody binding titers against the I53-50, I53-50A, and I3-01NS scaffolds. Each symbol represents an individual animal and the GMT from each group is indicated by a horizontal line. The dotted horizontal line represents the limit of detection for the assay. The dotted vertical line separates the protein and mRNA immunized groups. Statistical analyses have been omitted for clarity but can be found in **Supplementary Information. c,** Raw ELISA data and fits used to determine titers in (**b**). Each colored curve represents an individual animal.

**Extended Data Fig. 6.**
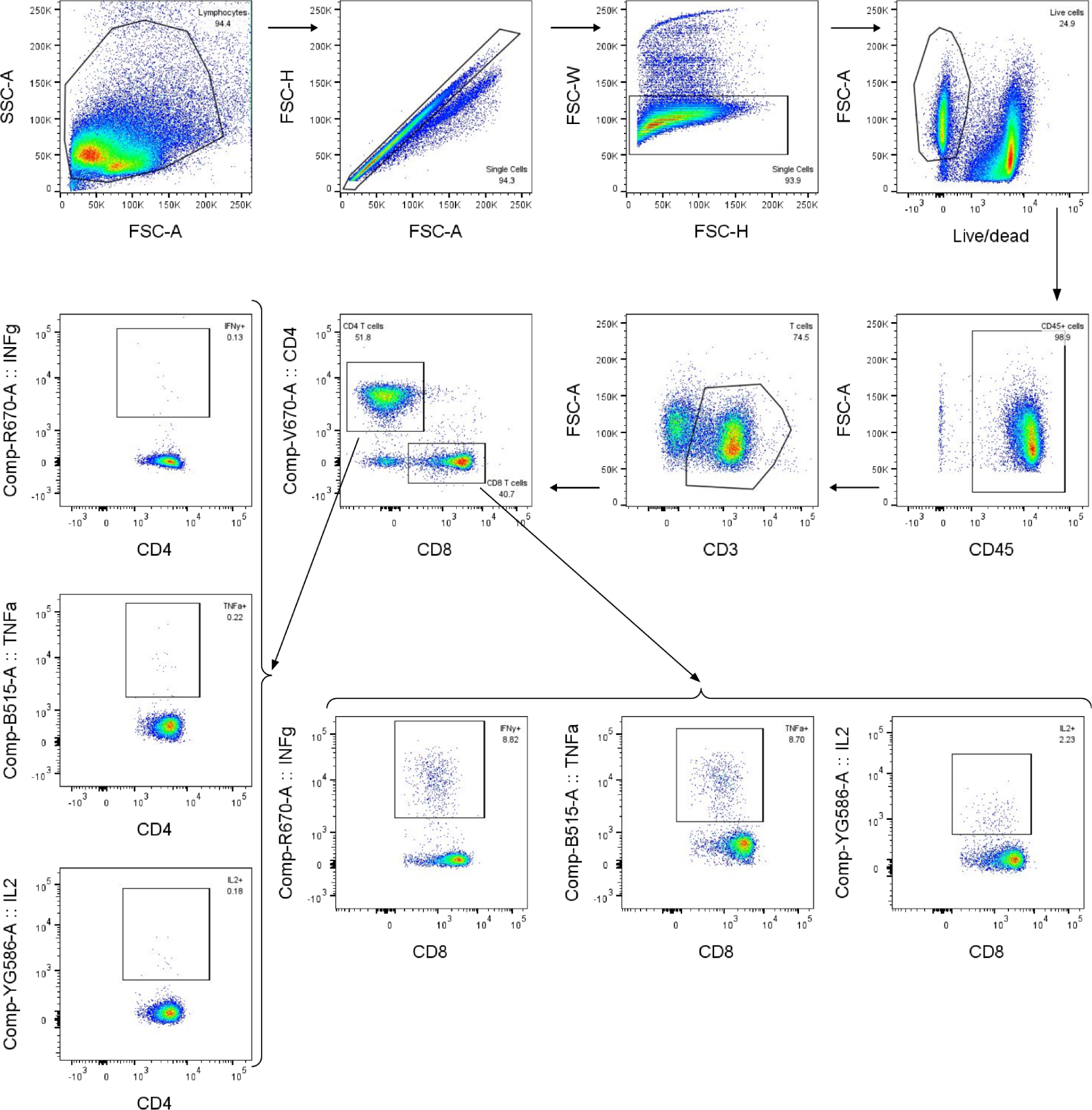
T cell gating strategy, related to Figure 3. Representative gating strategy for evaluating antigen-specific CD4 and CD8 T cells.

**Extended Data Fig. 7.**
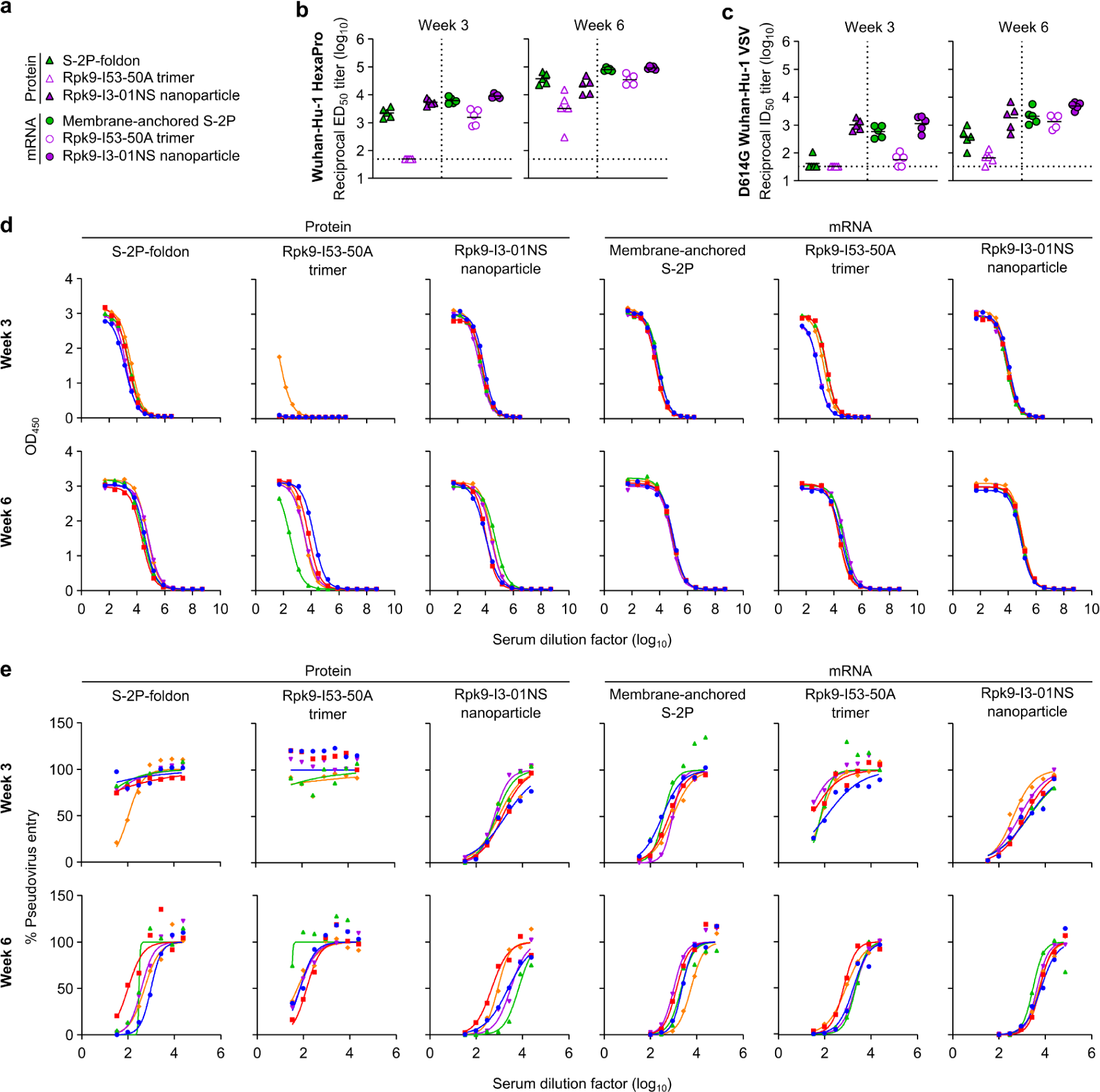
Antigen-specific and neutralizing antibody titers in C57BL/6 mice. **a,** Groups assessed for antigen-specific and neutralizing antibody responses (from mice in Fig. 3). **b,** Serum antibody binding titers against Wuhan-Hu-1 SARS-CoV-2 S HexaPro, measured by ELISA. **c,** Serum neutralizing antibody titers against VSV pseudotyped with D614G Wuhan-Hu-1 SARS-CoV-2 S. **d,** Raw ELISA data and fits used to determine titers in (**b**). **e,** Raw D614G Wuhan-Hu-1 SARS-CoV-2 pseudovirus entry data and fits used to determine titers in (**c**). **b-c,** The GMT from each group is indicated by a horizontal line. The dotted horizontal line represents the lowest limit of detection among the plotted data, as the limits of detection vary between groups. Statistical analyses have been omitted for clarity but can be found in **Supplementary Information. d-e,** Each color represents an individual animal.

**Extended Data Fig. 8.**
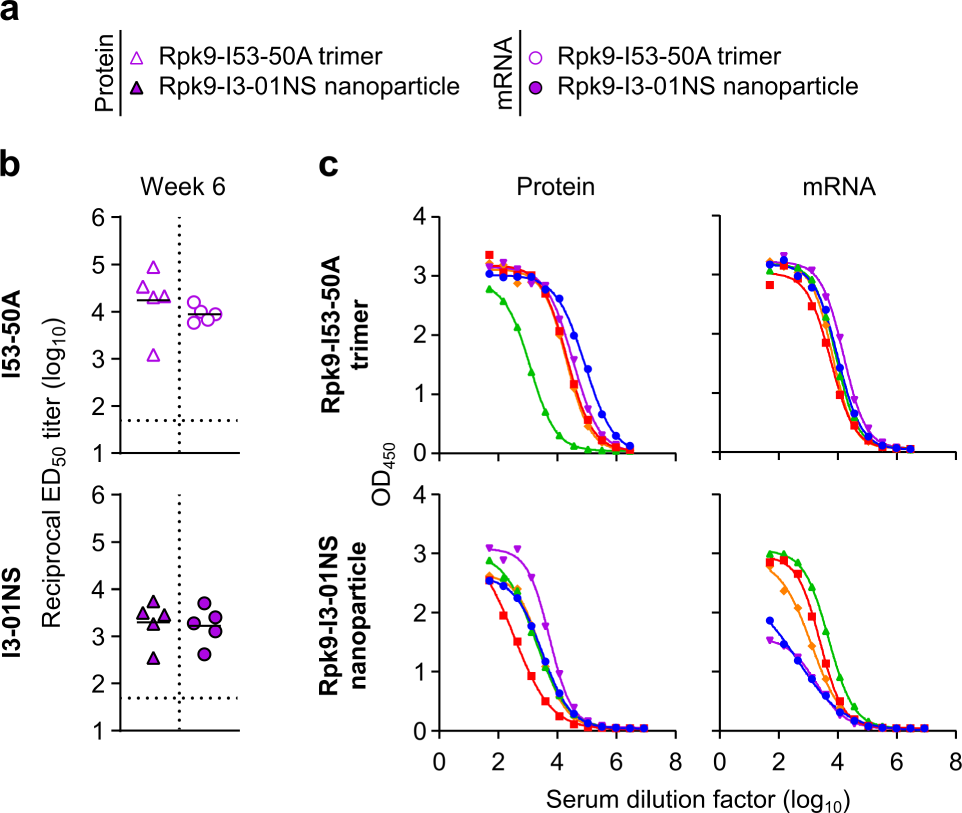
Anti-scaffold antibody titers in C57BL/6 mice three weeks post-boost. **a,** Groups assessed for anti-scaffold responses at week 6 (from mice in Fig. 3). **b,** Serum antibody binding titers against the I53-50A and I3-01NS scaffolds. Each symbol represents an individual animal and the GMT from each group is indicated by a horizontal line. The dotted horizontal line represents the limit of detection for the assay. Statistical analyses have been omitted for clarity but can be found in **Supplementary Information. c,** Raw ELISA data and fits used to determine titers in (**b**). Each color represents an individual animal.

**Extended Data Fig. 9.**
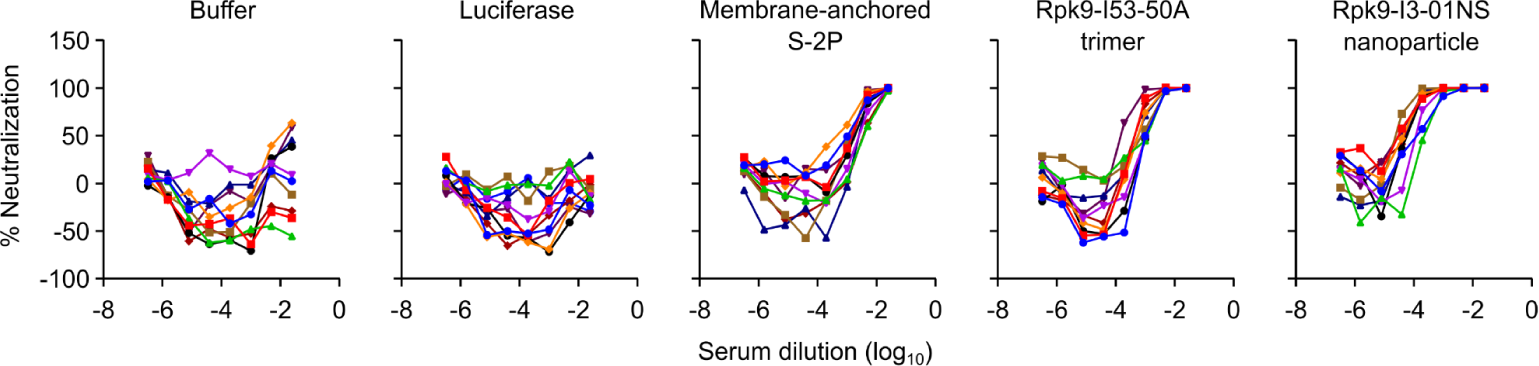
Raw data curves of D614G Wuhan-Hu-1 SARS-CoV-2 authentic virus neutralization data, related to Figure 4. Serum neutralizing activity against D614G Wuhan-Hu-1 SARS-CoV-2 authentic virus. Each color represents an individual animal. The mean of two technical replicates are shown.

**Extended Data Fig. 10.**
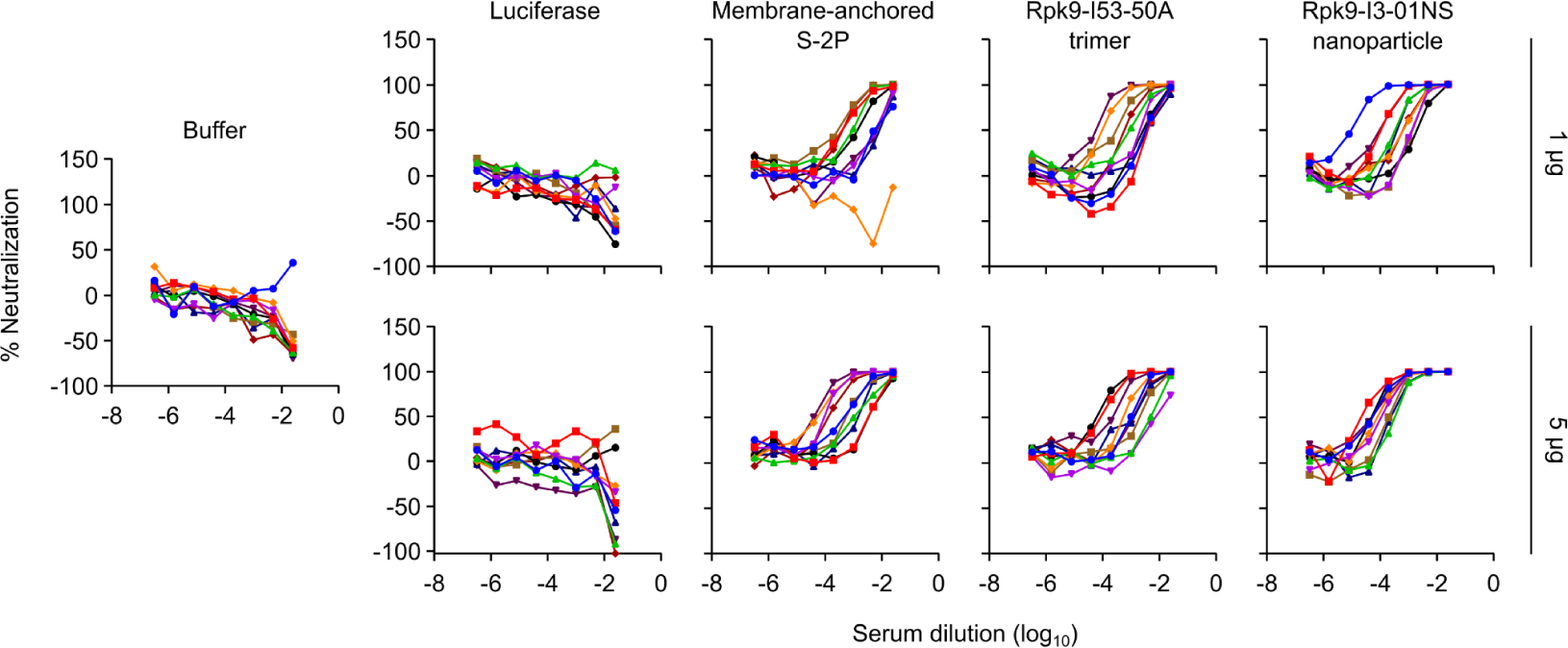
Raw data curves of Omicron BA.5 SARS-CoV-2 authentic virus neutralization data, related to Figure 5. Serum neutralizing activity against Omicron BA.5 authentic virus. Each color represents an individual animal. The mean of two technical replicates are shown.

**Supplementary Table 1.**
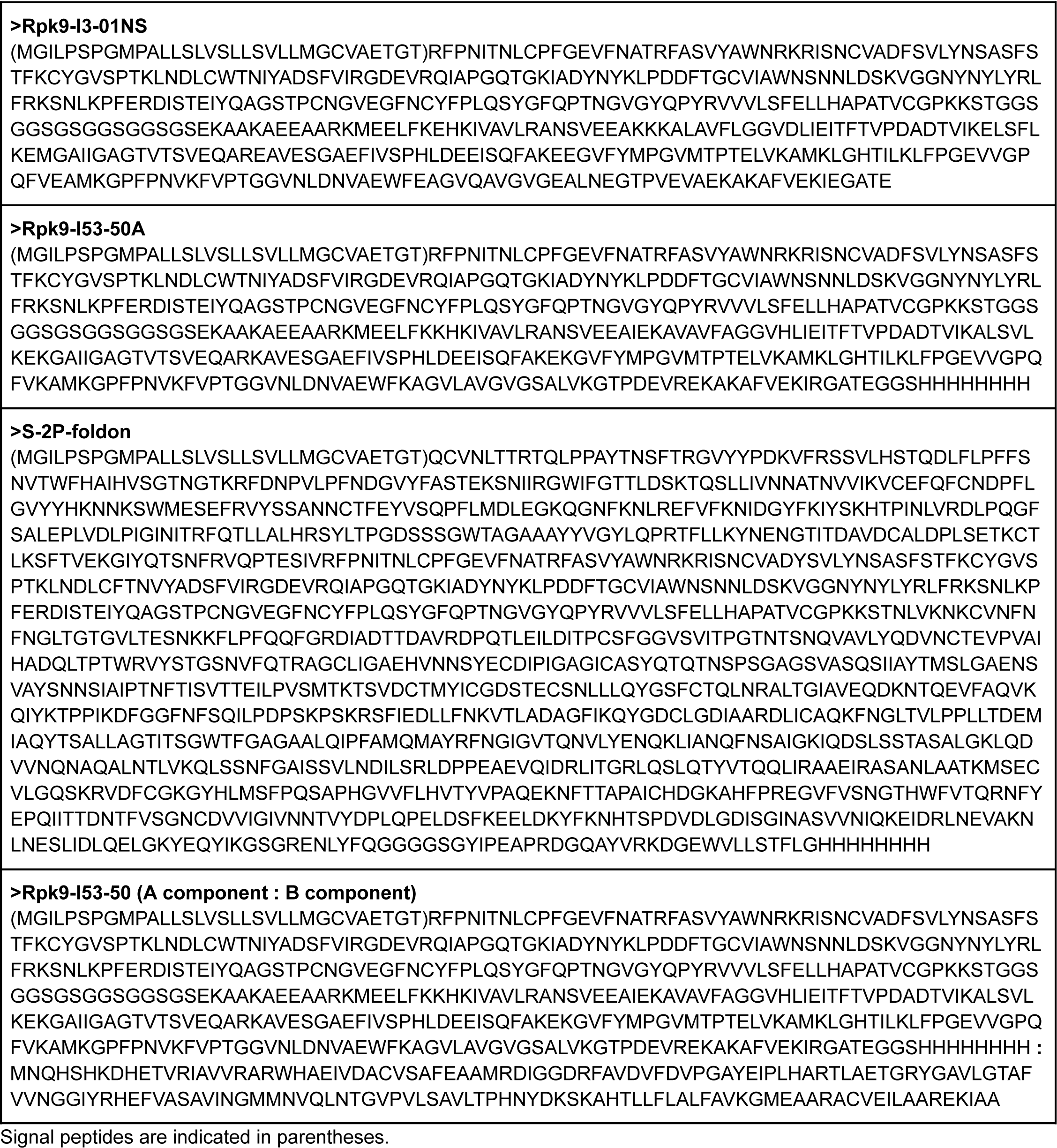
Amino Acid Sequences of Proteins Used for Immunizations.

**Supplementary Table 2.**
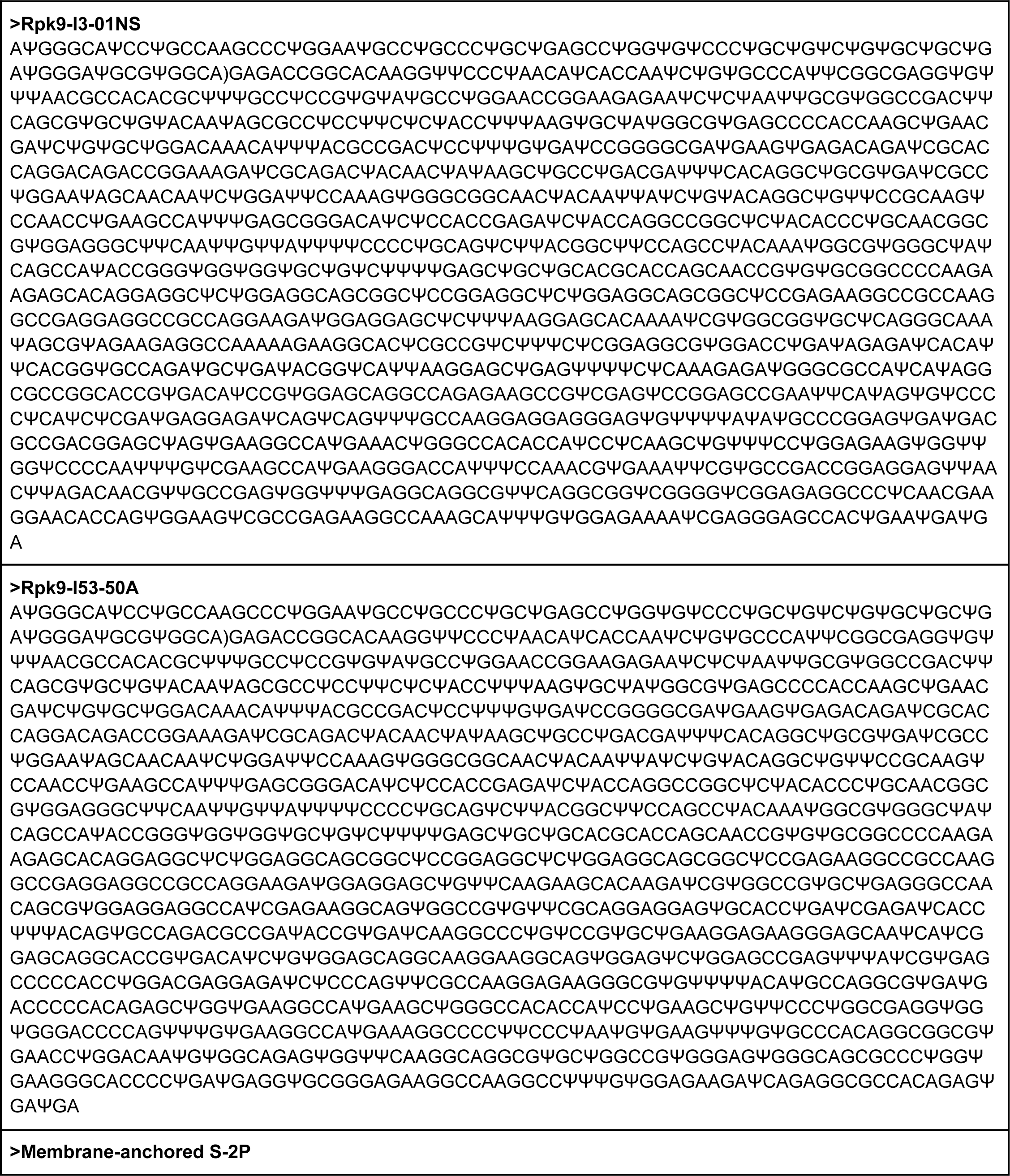
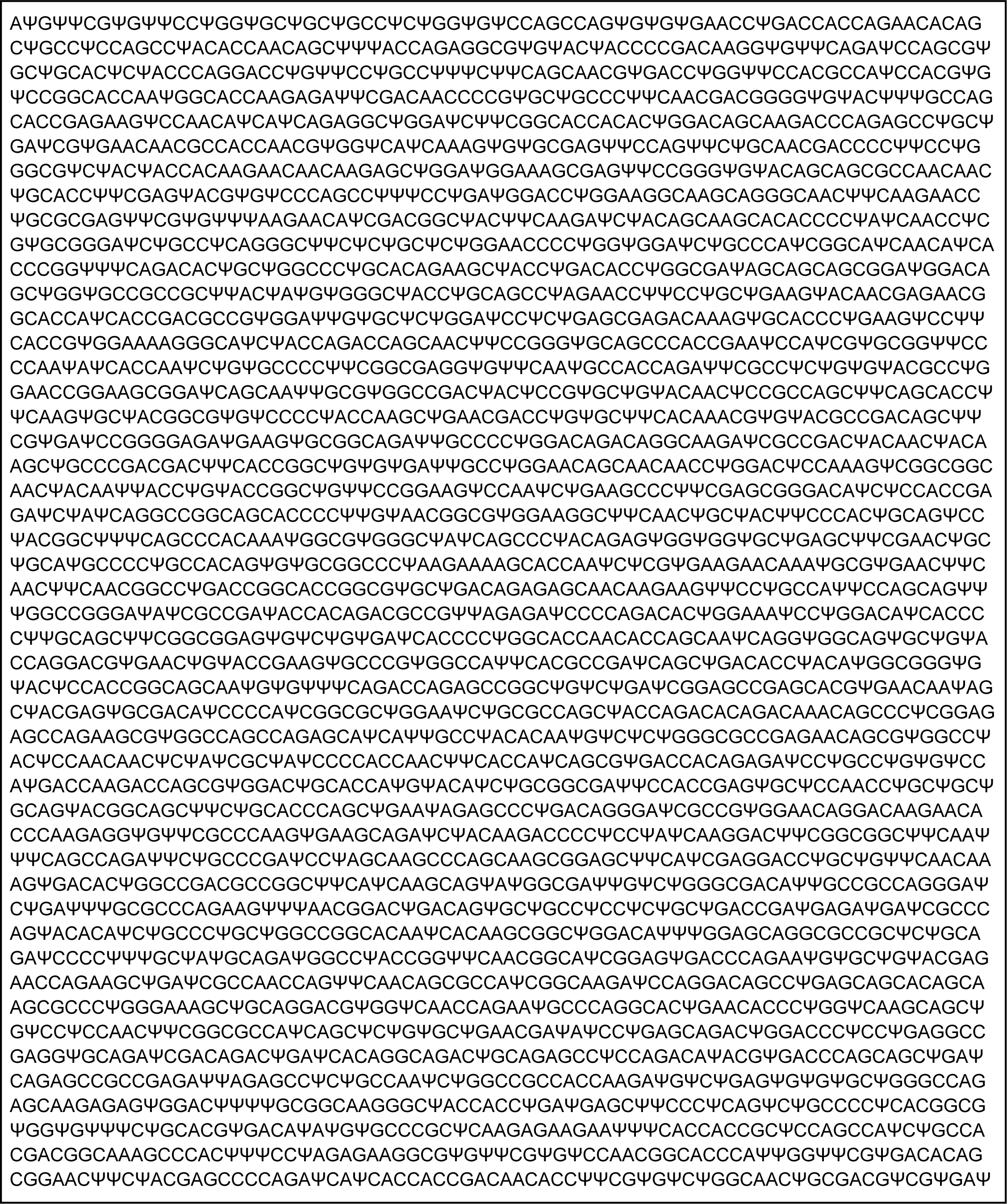
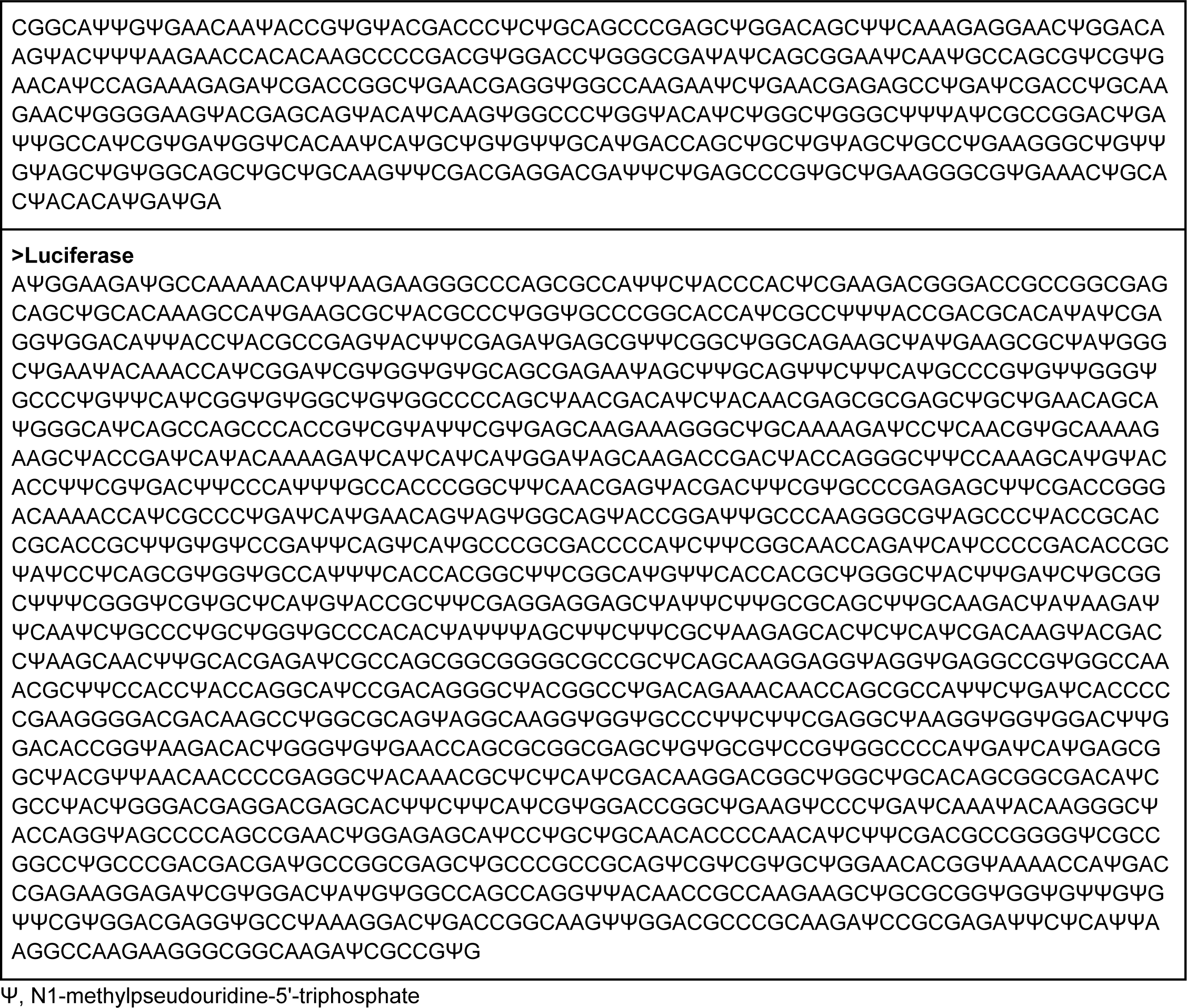
ORF Nucleic Acid Sequences of mRNA Constructs Used for Immunizations.

